# Graded optogenetic activation of the auditory pathway for hearing restoration

**DOI:** 10.1101/2022.09.05.506618

**Authors:** Artur Mittring, Tobias Moser, Antoine Huet

## Abstract

Optogenetic control of neural activity enables innovative approaches to improve functional restoration of diseased sensory and motor systems. For clinical translation to succeed, optogenetic stimulation needs to closely match the coding properties of the targeted neuronal population and employ suitable emitters at their optimal operation. This requires tailoring of channelrhodopsins, emitters and coding strategies. Here, we provide a framework to parametrize optogenetic neural control and apply it to the auditory pathway that requires high temporal fidelity of stimulation. We used viral gene transfer of the ultrafast targeting-optimized Chronos into spiral ganglion neurons (SGNs) of the cochlea. We characterized the light-evoked response by *in vivo* recordings from individual SGNs and neurons of the anteroventral cochlear nucleus (AVCN) that detect coincident SGN input. Our results demonstrate that spike probability of SGNs can be gradually dialed by adjusting the width of light pulses of constant intensity, which optimally serves efficient laser diode operation. We identified an effective pulse width of 1.6 ms to maximize information encoding in SGNs. An upper boundary of optical stimulation rates results from robust spike rate adaptation that required a few tens of milliseconds to recover. We developed a semi-stochastic stimulation paradigm to rapidly (within minutes) estimate the transfer function from light to SGNs firing. The semi-stochastic stimulus evoked firing of different statistics allowing to approximate the time constant of neuronal integration in the AVCN. Our data pave the way to design the sound coding strategies of future optical cochlear implants.

## Introduction

Optogenetics has revolutionized life sciences by enabling genetically, anatomically and temporally precise interrogation of organ function and of the brain and its neural circuits, in particular (for review, (Kim et al., 2017)). Optogenetics also unlocked new perspectives for restoring function in diseased sensory systems by bypassing dysfunctional or lost receptor cells via direct control of the activity of afferent neurons (for review, (Kleinlogel et al., 2020; Wolf et al., 2022)). Cellular specificity and spatial confinement of optogenetic stimulation promise an improvement of functional restoration beyond that amenable to electrical stimulation used in current cochlear and retinal implants. The clinical feasibility of this approach was recently proven by the first-in-human trial allowing partial vision restoration in a blind patient (Sahel et al., 2021). Similarly, optogenetics is an attractive alternative to electrical stimulation for restoring motor function or controlling chronic pain (Elkouzi et al., 2019; Gundelach et al., 2020; Kuner and Kuner, 2021)

Applying optogenetics to the first neurons of the auditory pathway, the spiral ganglion neurons (SGN), enables new opportunities for improved hearing restoration strategies (Dieter et al., 2020; Wolf et al., 2022). We recently demonstrated a major gain in spectral selectivity for optogenetic stimulation of the SGNs over the conventional electric stimulation used by electrical cochlear implants (eCI, (Dieter et al., 2020, 2019; Hernandez et al., 2014; Keppeler et al., 2020; Khurana et al., 2022)). This promises that future optical cochlear implants (oCI) will operate with a greater number of independent stimulation channels, which is expected to convey more natural sound perception and improve speech understanding in presence of background noise and music appreciation (Dieter et al., 2020; Wolf et al., 2022). However, improving the spectral code by optical stimulation should not trade in a poor temporal coding e.g. due to slow gating of channelrhodopsins (ChRs). Hence, efforts have been undertaken to speed up optogenetic control of SGNs by ultrafast ChRs (Bali et al., 2021; Keppeler et al., 2018; Mager et al., 2018).

The use case of optogenetic hearing restoration exemplifies that establishing optimal optogenetic control of neural activity in a given circuit is far from trivial. It requires a good tuning of membrane expression, matching the biophysical properties of the opsin (Klapoetke et al., 2014; Mager et al., 2018) to the neuron’s excitability (Herman et al., 2014), and identifying optimal simulation parameters also regarding the operation of the chosen emitters. For a neuronal population of interest, this typically necessitates empirical measurement of the transfer function from light to spike in different illumination conditions (Berndt et al., 2011; Herman et al., 2014; Klapoetke et al., 2014; Mattis et al., 2011). This is highly relevant also to the optogenetic stimulation of the auditory pathway. To date, the expression of various ChRs in SGNs has been achieved (channelrhodopsin-2: (Hernandez *et al*, 2014; Richardson *et al*, 2021); CatCh: (Dieter et al., 2019; Keppeler et al., 2020, p. 20; Wrobel et al., 2018); fast and very-fast-Chrimson variants: (Bali et al., 2021; Mager et al., 2018); and Chronos (Duarte et al., 2018; Keppeler et al., 2018)). In response to saturating light pulses delivered at a low repetition rate (10 -50 Hz), SGNs fire one (CatCh, Chronos and f- and vf-Chrimson) to 3 (CatCh) action potentials with a sub-millisecond precision (first spike jitter ∼ 0.25 ms,(Bali et al., 2021; Keppeler et al., 2018; Mager et al., 2018; Wrobel et al., 2018)).

Recordings from hippocampal cultured neurons showed that the maximum frequency at which light pulses entrain action potentials is co-determined by the photocurrent amplitude, the closing kinetics of the opsin, and the intrinsic maximum firing rate of the neuron (Berndt et al., 2011; Gunaydin et al., 2010). SGNs recorded *in vivo*, show a similar relationship when expressing ChRs with closing time constants ≥ 3 ms (at body temperature, (CatCh: Wrobel et al, 2018; f-Chrimson: Mager et al, 2018). Yet, the greatest temporal fidelity that was achievable with faster closing opsins such as Chronos (≤ 1ms) and vf-Chrimson (∼1.5 ms) did not enable light-synchronized firing to surpass 200 Hz of stimulation (Keppeler et al, 2018; Bali et al, 2021) suggesting an influence of other parameters than closing kinetics. Another important dimension for bionic coding, is the range of stimulus intensity represented by the neurons spike rate and temporal code. Recently, we showed a dynamic range of ∼ 4 or 1 dB (mW) in SGNs expressing f- or vf-Chrimson, respectively (Bali et al, 2021) which indicates that the dynamic range depends on the ChR employed.

To date, multiple open questions remain to be addressed *en route* to the application of optogenetics for hearing restoration. How can we gradually activate neurons at high light irradiance as required for efficient operation of semiconductor lasers (i.e. strong and submillisecond driver current pulses). Is there an optimal pulse width for efficient intensity encoding? Does the temporal precision of optically evoked action potentials comply with the integration window of the second-order neurons (i.e. the anteroventral cochlear nucleus, AVCN, neurons)?

Here, we addressed these questions in mice by *in vivo* recordings from single SGNs expressing the ultra-fast trafficking optimized Chronos (Chronos-ES/TS, τ_off_ = 0.76 ms at body temperature (Keppeler et al., 2018)). We demonstrate that spike probability can be gradually dialed by adjusting the width of light pulses like what can be achieved with acoustic clicks of different sound pressure levels. We identified an optimal pulse width of 1.6 ms for maximizing the amount of light intensity information in the spike trains. Further, we identified that high-rate optical stimulation induce spike failure that likely was due to depolarization block and which recovers in a few tens of milliseconds. Our data revealed that the timing of optogenetically evoked SGN spikes varies little across iterations for a given neuron but substantially within an SGN population, again reminiscent of what is observed acoustically. Finally, a novel semi-stochastic stimulus allowed us to rapidly (within minutes) establish the transfer function from light to SGN firing and to generate firing of different statistics for approximating the time constant of neuronal integration in AVCN neurons.

## Results

To characterize the encoding of optogenetic cochlear stimulation by the first two neuronal populations of the auditory pathway, we performed stereotaxic *in vivo* single unit recordings of SGNs and AVCN neurons in response to acoustic (**Figure 1.A**) or optogenetic **Figure 1.B**) stimulation. For optogenetic stimulation, SGNs were transduced by postnatal AAV injection to express the fast-closing, trafficking-optimized Chronos (Chronos-ES/TS) under the control of the human synapsin promoter (Keppeler et al., 2018). The light was delivered in the cochlea via an optical fiber coupled to a blue laser λ = 473 nm. Out of 55 injected mice, 49 showed an optogenetic activation of the auditory pathway characterized by optogenetically evoked auditory brainstem responses (oABR, **figure S1.A-C**, P_1_-N_1_ amplitude at 35 mW = 2.00 ± 0.41 μV, threshold = 12.19 ± 1.59 mW, average ± 95% confidence interval). Across all AAV-injected cochleae, 64.14 ± 5.97 % of the SGNs were transduced (**figure S1**) and the lowest oABR thresholds were measured for the cochleae with the highest transduction rate (**figure S1.K**, correlation coefficient = -0.44, *p*-value = 0.00039).

**Figure 1:**
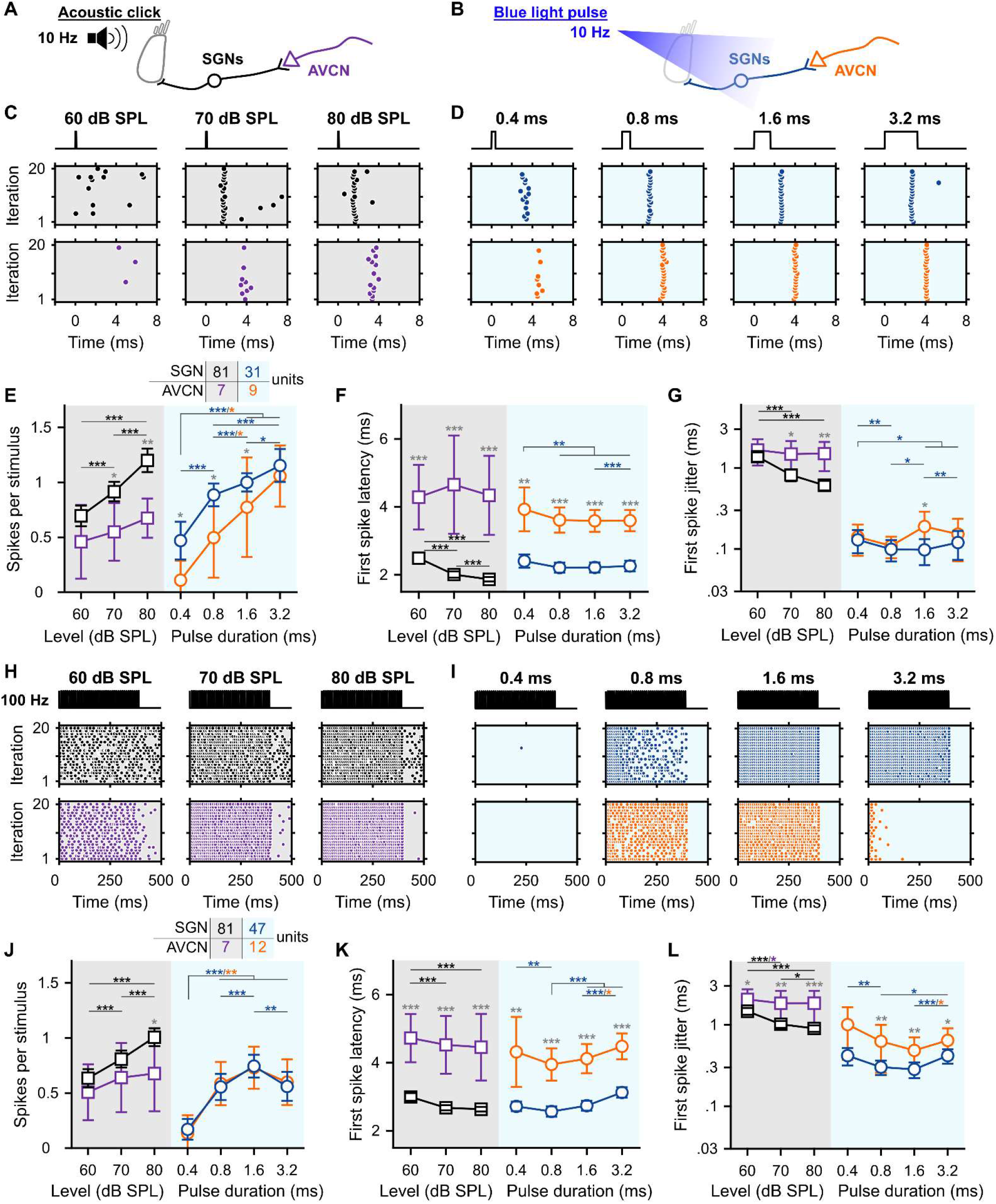
Gradual optogenetic activation of the first neurons of the auditory pathway with light pulses of increasing durations. **A-B**. Schematic of the *in vivo* single unit recordings using sharp micropipettes from spiral ganglion neurons (SGNs, black), optogenetically modified SGNs (SGNs, blue) or neurons of the anteroventral cochlear nucleus (AVCN, acoustic: purple and optogenetic: orange) in response to acoustic click (A) or optogenetic (B) stimulation delivered to the cochlea from an optical fiber (400 ms stimulation followed by 100 ms silence/dark, 10-20 trials per condition, all data are paired). **C-D**. Representative raster plots from one SGN (top panel) and one AVCN neuron (bottom panel) in response to single acoustic clicks at various sound pressure levels (C, click trains presented at 10 Hz) or single light pulses of different pulse durations (D, ∼35 mW light pulses delivered at 10 Hz). Note that acoustic and optogenetic experiments were done from two distinctive cohorts of animals, but SGNs and AVCN neurons were recorded from the same animals. **E-G**. Quantification of the number of spikes per stimulus (E), first spike latency (F) and first spike jitter (G) as a function of the click level for the acoustic modality (gray background) or light pulse duration for the optogenetic modality (blue background). The color code is similar to panels A-D. The number of units per group is presented in the inset of panel E. Data are represented as mean and confidence interval (95%). The effect of the acoustic levels and light pulse durations was tested by a Wilcoxon signed rank test on paired samples followed by a Bonferroni correction of the *p*-values. The difference between SGNs and AVCN neurons was tested by a two-sample t-test or a Wilcoxon rank sum test according to the outcome of Jarque-Bera normality testing (grey symbols; *, *p*-value ≤ 0.05; **, *p*-value ≤ 0.01; ***, *p*-value ≤ 0.001). **H-L**. Same as C-G in response to 100 Hz acoustic click or light pulse trains. Note that the units presented in H-I are the same as in C-D.

### Graded optogenetic activation of spiral ganglion neurons by light pulses of increasing duration

Studying the integration mechanisms of an already identified neural network would benefit from gradually activating it to generate various spike patterns and use them to measure the transfer function from one element of the circuit to another one. Gradual neuronal activation can be achieved optogenetically by controlling the number of photons reaching the channelrhodopsins (ChR) expressed at the membrane of neurons. Experimentally, this can be done by varying the radiant flux and/or the light pulse duration (Berndt et al., 2011, 2011; Keppeler et al., 2018; Klapoetke et al., 2014; Mager et al., 2018; Wrobel et al., 2018). The second has the advantage of not being associated with a spatial change of the light cone (Khurana et al., 2022). Additionally, laser diodes, candidate semiconductor emitters for future optical cochlear implants, show best efficiency for strong and brief driver currents which would also require gradual SGN activation with different light pulse durations (Rogers, 2021).

First, we assessed the encoding of optogenetic and acoustic stimulus strength in SGNs. We varied light pulse duration at fixed strong intensity (radiant flux). We compared optogenetic responses to those evoked by brief acoustic stimuli (submillisecond clicks of variable intensity but fixed duration) recorded from non-optogenetically modified SGNs (**figure 1.C-D**, top row; optic, *n* = 31 SGNs recorded from 6 mice, maximal laser output stimulation = 35 ± 0.35 mW, average ± 95% confidence interval; acoustic, *n* = 81 SGNs recorded from 8 mice). All optogenetic or acoustic conditions were recorded pairwise and therefore can be directly compared per neuron. The number of spikes per stimulus increased with the light pulse duration from 0.47 ± 0.17 at 0.4 ms to 1.15 ± 0.15 at 3.2 ms (*p*-value = 1.03 × 10^−5^, Wilcoxon signed rank test on paired samples followed by a multi-comparison test) ; and with the acoustic click intensity (**figure 1.C-E**) from 0.69 ± 0.09 at 60 dB SPL_PE_ to 1.2 ± 0.11 at 80 dB SPL_PE_ (*p*-value = 2.84 × 10^−14^). The optimal representation of ∼1 spike per stimulus was achieved by 1.6 ms light pulse and acoustic click between 70 and 80 dB SPL. Both, varying light pulse duration and click intensity gradually activated the input of the auditory pathway.

The latency and temporal precision of the first spike elicited per stimulus also showed a strong dependence on the stimulation strength (**figure 1.F-G**). Optogenetically, the first spike latency was significantly longer at 0.4 ms (2.40 ± 0.20) than at 0.8, 1.6 and 3.2 ms (2.21 ± 0.15, 2.22 ± 0.15 and 2.26 ± 0.16 respectively, *p*-values ≤ 0.0021, Wilcoxon signed rank test on paired samples followed by a Bonferroni correction). Temporal precision was greatest at 1.6 ms (first spike latency jitter = 0.10 ± 0.03 ms respectively) than 0.4, 0.8 and 3.2 ms (0.13 ± 0.04, 0.10 ± 0.03 and 0.12 ± 0.05 ms respectively, *p*-values ≤ 0.0151). Acoustically, the first spike latency decreased exponentially from 2.49 ± 0.17 ms at 60 dB SPL_PE_ to 1.86 ± 0.06 ms at 80 dB SPL_PE_ (*p*-value = 8.19 × 10^−11^). Likewise, the jitter of first spike latency decreased from 1.37 ± 0.18 ms at 60 dB SPL_PE_ to 0.61 ± 0.08 ms at 80 dB SPL_PE_ (*p*-value = 5.51 × 10^−9^). Surprisingly, given the need for cochlear sound processing upstream of SGN spike generation, the first spike latency in response to 80 dB SPL_PE_ acoustic click was significantly shorter than all optogenetically tested conditions (*p*-values ≤ 0.0019, Kruskal-Wallis test followed by a Tukey’s Honestly Significant Difference Procedure). Yet, direct light activation of SGNs showed smaller (one order of magnitude) temporal jitter of the first spike for optogenetic compared to acoustic stimulation.

Aside from SGNs, we also recorded from AVCN neurons (see methods section for differentiation between recordings from SGNs and AVCN neurons). We discarded about 30% of the AVCN neurons that encoded the stimulus weakly (< 0.3 spikes/stimulus for any of the tested conditions). For all conditions tested, the first spike latency of AVCN neurons was significantly longer than for SGNs reflecting the SGN conduction and synaptic transmission (**figure 1.C, F**, *p*-values ≤ 0.0033, Jarque-Bera test followed accordingly by a two-sample t-test or a Wilcoxon rank sum test). For most tested conditions, the representation of the stimulus in terms of the number of spikes was weaker for AVCN neurons compared to SGNs (0.4, 0.8 and 1.6 ms light pulse: *p*-values ≤ 0.0159; 70 and 80 dB SPL_PE_ acoustic clicks: *p*-values ≤ 0.0114). Interestingly, this difference decreased for optogenetic stimulation with increasing stimulation duration (strength) from 0.36 at 0.4 ms to 0.09 at 3.2 ms. For acoustic stimulation, in contrast, the difference increased with stimulation strength from 0.24 at 60 dB SPL_PE_ to 0.53 at 80 dB SPL_PE_ suggesting a better recoding of optogenetic stimuli compared to acoustic stimuli by the recorded AVCN neurons.

Next, we investigated if the graded activation of the auditory pathway by various duration of individual light pulses was present also during light pulse trains of high repetition rates (100, 316 and 1000 Hz, **figure 1.H-L, figure S2**). Again, we compared optogenetic stimulation to acoustic stimulation, here embarking on click trains presented at various intensities. At 316 and 1000 Hz, the firing of most SGNs fully adapted (i.e. spike rate decreased to 0 spikes/s) within the first few presentations of the light pulses (**figure S2.C-F**). This observation seemed unexpected given the fast closing kinetics reported for Chronos (Keppeler et al., 2018; Klapoetke et al., 2014). However, such rapid spike rate adaptation is consistent with our previous recordings of SGN firing mediated by Chronos-ES/TS (Keppeler et al., 2018) and vf-Chrimson (Bali et al., 2021). Hence, we focused the analysis on 100 Hz stimulation trains. While the number of spikes per stimulus was generally lower for light pulse trains, we observed a dependence of SGN firing on stimulus strength similar to that found for individual light pulses/clicks. Optogenetically the best representation of the light pulse in the SGNs firing was achieved at 1.6 ms (0.74 ± 0.10 spike/stimulus) and decreased for shorter and longer pulse durations (0.4 ms: 0.17± 0.09; 0.8 ms: 0.56 ± 0.12; 3.2 ms: 0.56 ± 0.13, *p*-values ≤ 0.0012, Wilcoxon signed rank test on paired samples followed by a Bonferroni correction). Spike rate adaptation (i.e. adaptation ratio ≥ 1, the ratio of the discharge rate between the first 50 ms of the train and the first 400 ms) was evident for all pulse durations. Adaptation was significantly less at 1.6 ms compared to 0.8 and 3.2 ms (**figure S2.B**, p-values ranging between 0.001 and 0.0012, Wilcoxon signed rank test on paired samples followed by a Bonferroni correction). Upon acoustic stimulation, the number of spikes per stimulus increased linearly with the sound level from 0.63 ± 0.08 at 60 dB SPL_PE_ to 1.00 ± 0.08 at 80 dB SPL_PE_. Adaptation of the firing rate was not observed at any level (adaptation ratio ∼ 1, **figure S2.B**). The timing and temporal precision of the first spike were differently affected by the stimulation strength between the 2 modalities. Optogenetically, the shortest first spike latency was measured at 0.8 ms (**figure 1.K**, 2.56 ± 0.15 ms, *p*-values ≤ 0.0093) and precision was highest for 0.8 and 1.6 ms (**figure 1.L**, first spike jitter = 0.30 ± 0.06 ms and 0.28 ± 0.06 ms respectively, *p*-values ≤ 0.02 for comparison to 0.4 and 3.2 ms). Acoustically, first spike latency and jitter decreased exponentially with the sound level. It is worth mentioning that acoustically, the temporal precision (i.e. the inverse of the spike jitter) was similar for single pulses and 100 Hz train stimuli, while optogenetically the precision decreased by a factor of ∼3.

At the AVCN level, the number of spikes per light pulse was comparable to the SGNs (**figure 1.J**). In contrast, the number of spikes per acoustic click was reduced in the AVCN neurons compared to the SGNs (80 dB SPL_PE_, *p*-value = 0.0257). For all modalities and conditions, the first spike precision was lower for AVCN neurons than SGNs (**figure 1.L**, p-values ≤ 0.0268) and, in general, was less sensitive to the stimulation strength.

Encoding of light pulses varied widely across SGNs, which was also found for SGNs recorded from the same animal and seemed independent of the transduction rate measured in their cochleae (**Figure 2.A-C**, *n* = 35 SGNs from 16 mice, 100 Hz light pulse train of 1.6 ms, correlation coefficient). The origin of this heterogeneity is currently unknown. Candidate mechanisms include heterogeneous ChR expression by the SGNs, different distances to the light source or varying neuron excitability. As a result, AVCN neurons might receive excitatory inputs that are less synchronous than originally quantified from individual optogenetically driven SGNs. To approximate this effect, we computationally simulated a population of SGNs from our SGN recordings and measured the statistics of the spike trains across SGNs per stimulus presentation (**figure 2.D-F**, see Methods for details, 50 bootstraps per condition; optogenetics, *n* = 47 SGNs; acoustic, *n* = 20 SGNs with the best frequency within the 16 kHz octave band). As expected, the number of spikes per light pulse (**figure 2.D**) and the latency of the first spike (**figure 2.E**) quantified at the population level were similar for all conditions to the average of the single SGNs. In contrast, the first spike latency jitter was systematically higher at the population level than within single SGNs and amounted to half of the population spike jitter measured acoustically (**figure 2.F**). This suggests that the responses to acoustic and optogenetic modalities are not as dissimilar as suggested by the quantification made on single neurons.

**Figure 2:**
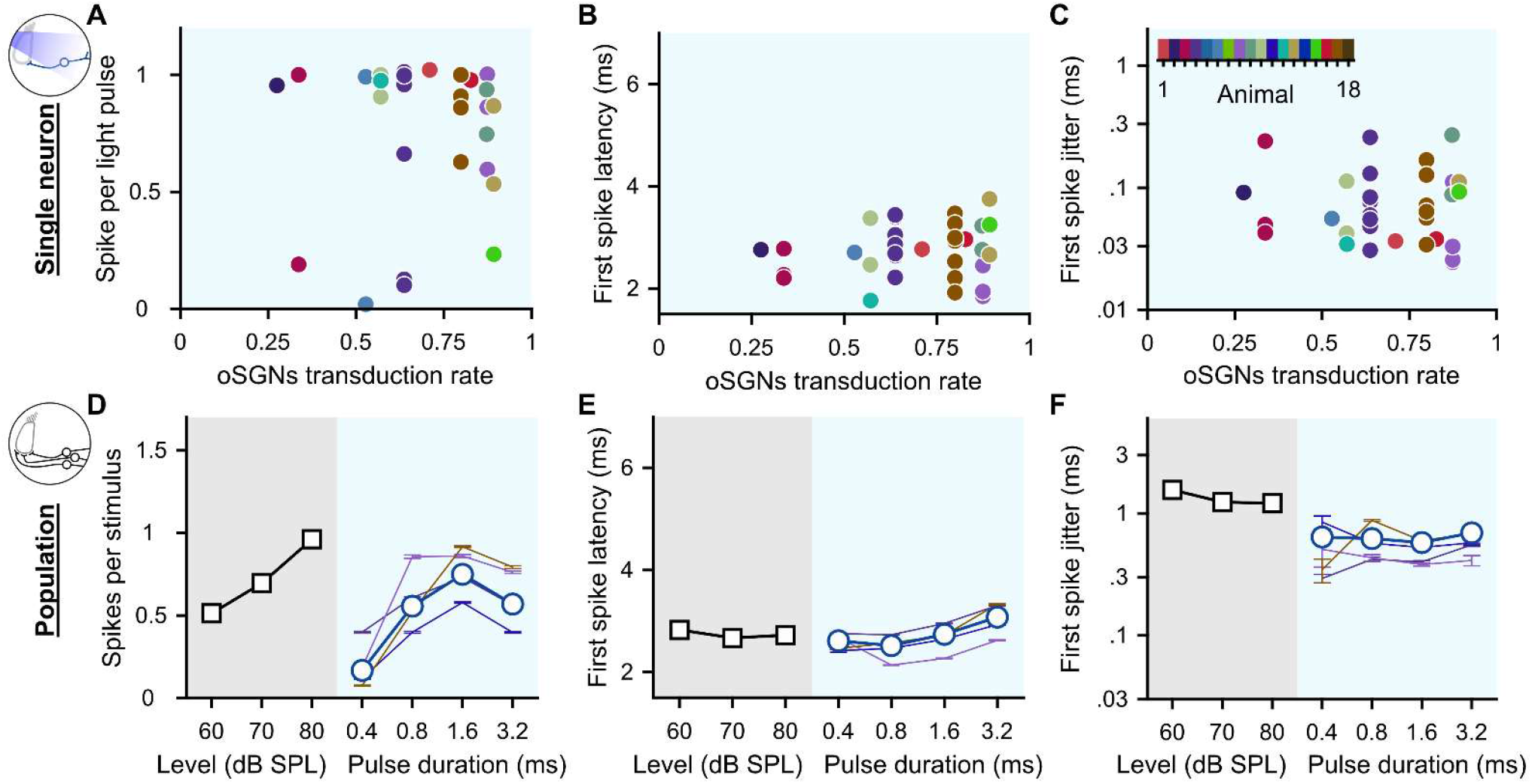
Heterogeneous optogenetic activation of the SGNs and artificial population response. **A-C**. Quantification of the spike per light pulse (A), first spike latency (B) and first spike jitter (C) as a function of the average SGNs transduction rate. Measurements were done in response to 100 Hz light pulse trains of 1.6 ms (*n* = 35 SGNs). A color code was used per animal. No correlation was found (correlation coefficient test). **D-F**. Quantification of the number of spikes per stimulus (D), first spike latency (E) and first spike jitter (F) computed across SGN recorded from a single animal (≥ 4 SGNs, thin colored line, same color code as in A-C) and all recorded SGNs (grey background, acoustic: *n* = 20 SGNs with the best frequency within the 16 kHz octave band; blue background, optogenetics: *n* = 47 SGNs).

Together our results show that varying the duration of high radiant flux light pulses enables a graded activation of SGNs similarly to changing the intensity of acoustic click. The best SGN encoding was obtained with 1.6 ms pulses and those spike trains were characterized by a high spike time precision (compared to acoustic stimulation). Contrasting with previous studies using acoustic stimulus on cats and Mongolian gerbils (Joris et al., 1994; Wei et al., 2017), no enhancement of the spike rate or temporal precision was observed within the small population of AVCN neurons recorded compared to the SGNs, which applied to both acoustic and optogenetic stimulation.

### Identifying the optimal pulse duration to encode light pulse amplitude

Next, we covaried pulse duration and light pulse amplitude (radiant flux) for a more comprehensive analysis of optogenetic coding. Acoustically, the stimulus intensity is encoded by individual SGNs by changes in the firing rate over ∼5 – 50 dB with different activation thresholds (Huet et al., 2016; Liberman, 1978; Sachs and Abbas, 1974; Taberner and Liberman, 2005; Winter et al., 1990)). Hence, each of them encodes intensity in a fraction of the audible dynamic range of ∼120 dB. The response of the SGNs to sound of various intensities is shaped by multiple mechanisms such as active micromechanics (Yates et al., 1990) and inner hair cell synapses (Ohn et al., 2016; Özçete and Moser, 2021), which are bypassed in direct SGN stimulation by cochlear implants. Using electric pulses the dynamic range per SGNs is limited to ∼1 dB (current level) (Miller et al., 2006). We recently reported intensity encoding by ChR-expressing SGNs and the measured dynamic ranges were dependent on the opsin used ranging from ∼1 and ∼3.5 dB (mW) (see (Bali et al., 2021) for discussion of dB (mW)).

To address optogenetic stimulus intensity coding, we recorded pairwise the response to 100 Hz light pulse trains of 0.4, 0.8, 1.6 and 3.2 ms at as many radiant fluxes as amenable within the time the recording lasted (**figure 3.A**, 20 iterations, 400 ms light on, 100 ms dark). A total of 29 SGNs from 12 mice were included for which at least 4 radiant fluxes were presented and where for at least one of the light pulse duration we covered the dynamic range of the SGNs from below its threshold to above its saturation (see Methods for details). To facilitate comparison between animals, radiant fluxes were converted to a level relative to the oABR threshold (as introduced in (Bali et al., 2021), see Methods, dB_rel to oABR thr._). At the threshold, the optogenetic SGN response is characterized by a strong adaptation of firing during the first ∼100 ms (Bali et al., 2021) following stimulus onset, therefore rate-level functions (RLF, **figure 3.B**) were built from the adapted rate (between 100 and 400 ms, green box in **figure 3.A**).

**Figure 3:**
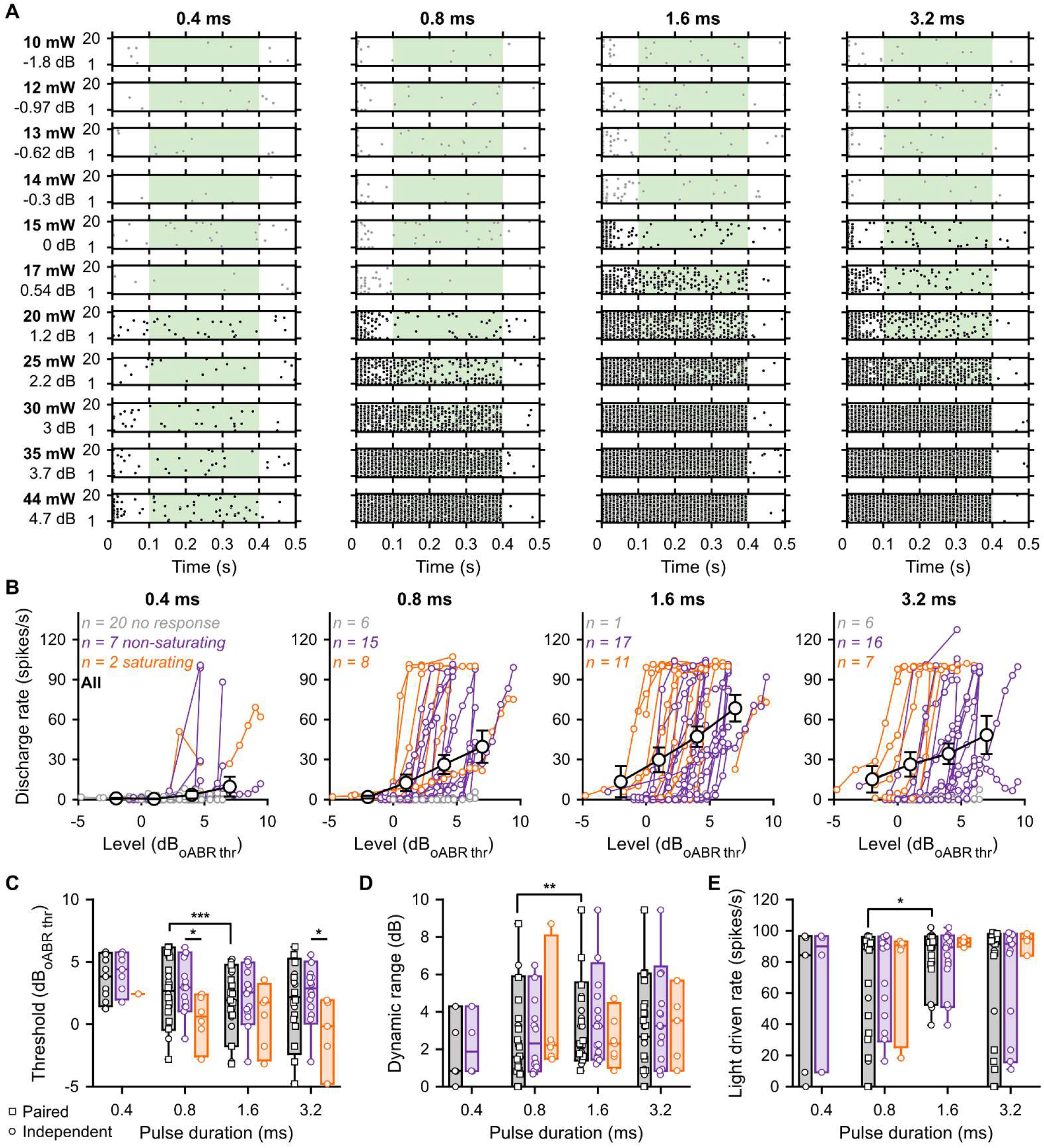
The best intensity encoding is achieved using 1.6 ms light pulses. **A**. Raster plots of a representative SGN evoked by 100 Hz light pulse trains (400 ms light, 100 ms dark) at various light intensities and pulse durations. The tested light intensities on the left are expressed in radiant flux (mW) and level relative to the oABR threshold (dB_oABR thr._, see Methods). Spike trains were computed on the adapted rate (100 to 400 ms, green window). Black dots highlight response above the threshold which was determined as the light intensities for which d’ ≥ 1 (see Methods). **B**. Rate level function of 29 SGNs measured in response to light pulse trains using various light pulse durations. A color code was used to represent non-responding (gray), non-saturating (purple) and saturating (orange) units (see Methods for classification). The average (± 95% confidence interval) rate-level function is plotted in black for all pulse durations (intensities values binned between -5 and 8 dB, bin width = 3 dB). Note that the highest light-driven rate is obtained at 1.6 ms. **C-E**. Quantification of the threshold (C), dynamic range (D) and light-driven rate (E) as a function of the light pulse duration. The same color code is similar to in B. Box plots show minimum, 25th percentile, median, 75th percentile, and maximum with individual data points overlaid. Circles represent independent data points and squares paired data points at 0.8, 1.6 and 3.2 ms. The effect of light pulse durations was tested at 0.8, 1.6 and 3.2 ms using a Wilcoxon signed rank test on paired samples followed by a Bonferroni correction. Following assessment for normality using a Jarque-Bera test, the difference between non-saturated and saturated units was tested accordingly by one-way analysis of variance or a Kruskal-Wallis test (*, p-value ≤ 0.05; **, p-value ≤ 0.01; ***, p-value ≤ 0.001).

RLFs classified as non-responding, saturating, or non-saturating were found for every tested pulse duration with the highest proportion of responding units observed at 1.6 ms (28/29 vs. 9, 23 and 23 at 0.4, 0.8 and 3.2 ms respectively, **figure 3.B**). When possible, a rate-based threshold was determined using the detection theory (see Methods, (Bali et al., 2021; Huet et al., 2018; Macmillan and Creelman, 2004)). The threshold tended to decrease with the pulse duration. Compared to 0.8 ms light pulses (2.87 ± 1.27 dB_rel oABR thr._, *n* = 19 SGNs, **figure 3.C**), the lowest threshold per SGN were measured at 1.6 ms (1.50 ± 1.12 dB_rel oABR thr._, *p*-value = 0.0006, Wilcoxon signed rank test on paired samples followed by a Bonferroni correction). It is worth noting that at 0.8 and 3.2 ms, the RLFs classified as saturated were activated ∼2.5 dB lower than non-saturated RLFs (p-values ≤ 0.0204, Jarque-Bera test followed accordingly by one-way analysis of variance or a Kruskal-Wallis test). Next, we measured the dynamic range (i.e. level difference from threshold to 99% of the maximum adapted firing rate for saturating SGN or to the highest tested level for non-saturating SGN, *n* = 21 SGNs, **figure 3.D**) and light-driven rate range (i.e. discharge rate difference from threshold to 99% of the maximum driven rate for saturating SGN or to highest tested level for non-saturating SGN, *n* = 21 SGNs, **figure 3.E**). Again 1.6 ms stimulation generated the widest dynamic and driven rate ranges (3.26 ± 0.96 dB and 84.45 ± 7.61 spikes/s) compared to 0.8 (2.41 ± 1.08 dB, *p*-value = 0.0084; 58.52 ± 18.07 spikes/s, *p*-value = 0.0191). These results suggest that the best intensity encoding in terms of the number of recruited SGNs, threshold, dynamic and discharge rate ranges was achieved using 1.6 ms light pulse duration.

### Recovery from light-induced refractoriness

In response to ongoing intense sounds (e.g. a few tens of minutes), SGNs continue to fire at a few hundred spikes per second (Kiang et al., 1965). Optogenetically, most SGNs tonically encode light pulse trains up to a stimulation rate of one to four hundred Hertz (which depends on the biophysical properties of the opsin) and fully adapted at higher stimulation rates (**figure S2**, (Bali et al., 2021; Keppeler et al., 2018; Mager et al., 2018; Wrobel et al., 2018)). How long does it take SGNs to recover from refractoriness and be responsive again?

To address this question, we designed a so-called forward masking protocol (**figure 4.A**, 20 iterations) composed of a masker (10 light pulse of 1.6 ms presented at 316 Hz) followed by a single light pulse (1.6 ms) presented at different time intervals (Δ t) ranging from 4 to 180 ms. We primarily employed the maximal radiant flux of our system (35 ± 0.35 mW) and then repeated the experiment at as many intensities as possible to assess if the recovery was level dependent. We recorded a total of 59 SGNs from 11 mice. In about 15% of recorded SGNs, the firing was not adapted by the masker (**figure 4.C**) and therefore recovery could not be assessed. For the adapted unit, a normalized spike probability curve was built (**figure 4.B, D**) and fitted by a mathematical model to extract the time of absolute refractoriness (i.e. the time needed to recover any firing, see Methods) and relative recovery (i.e. the time needed to recover 95% of the normalized spike probability). The absolute refractoriness was level independent and amounted to 7.4 ± 0.67, 16.45 ± 3.88 and 12.98 ± 2.64 ms at 3, 5 and 7 dB_rel. to oABR thr._ (**figure 4.E**, average ± 95% confidence interval, bin width = 2 dB, Jarque-Bera test followed by a Kruskal-Wallis test). Similarly, the time to relative recovery was level independent and amounted to 36.56 ± 18.98, 44.01 ± 17.60 and 27.61 ± 9.41 ms at 3, 5 and 7 dB_rel. to oABR thr_ (**figure 4.F**). Recovery time varied starkly and spanned over two orders of magnitude, thus revealing another major heterogeneity among SGNs. Acoustically, the time of recovery from a previous stimulus depends on the spontaneous activity of the given SGN: SGNs with a spontaneous activity ≥ 18 spikes/s recover the fastest (Relkin and Doucet, 1991). We did not observe a correlation between the time to relative recovery and the spontaneous activity (**figure S3.A**) or the number of spikes elicited per light pulse (**figure S3.B**) in our data. We note, that the spontaneous activity of SGNs was highly reduced following the cochleostomy and the insertion of the optical fiber compared to physiological conditions (Taberner and Liberman, 2005) and amounted to 0.29 ± 0.11 spikes/s with the highest value at 2.04 spikes/s.

**Figure 4:**
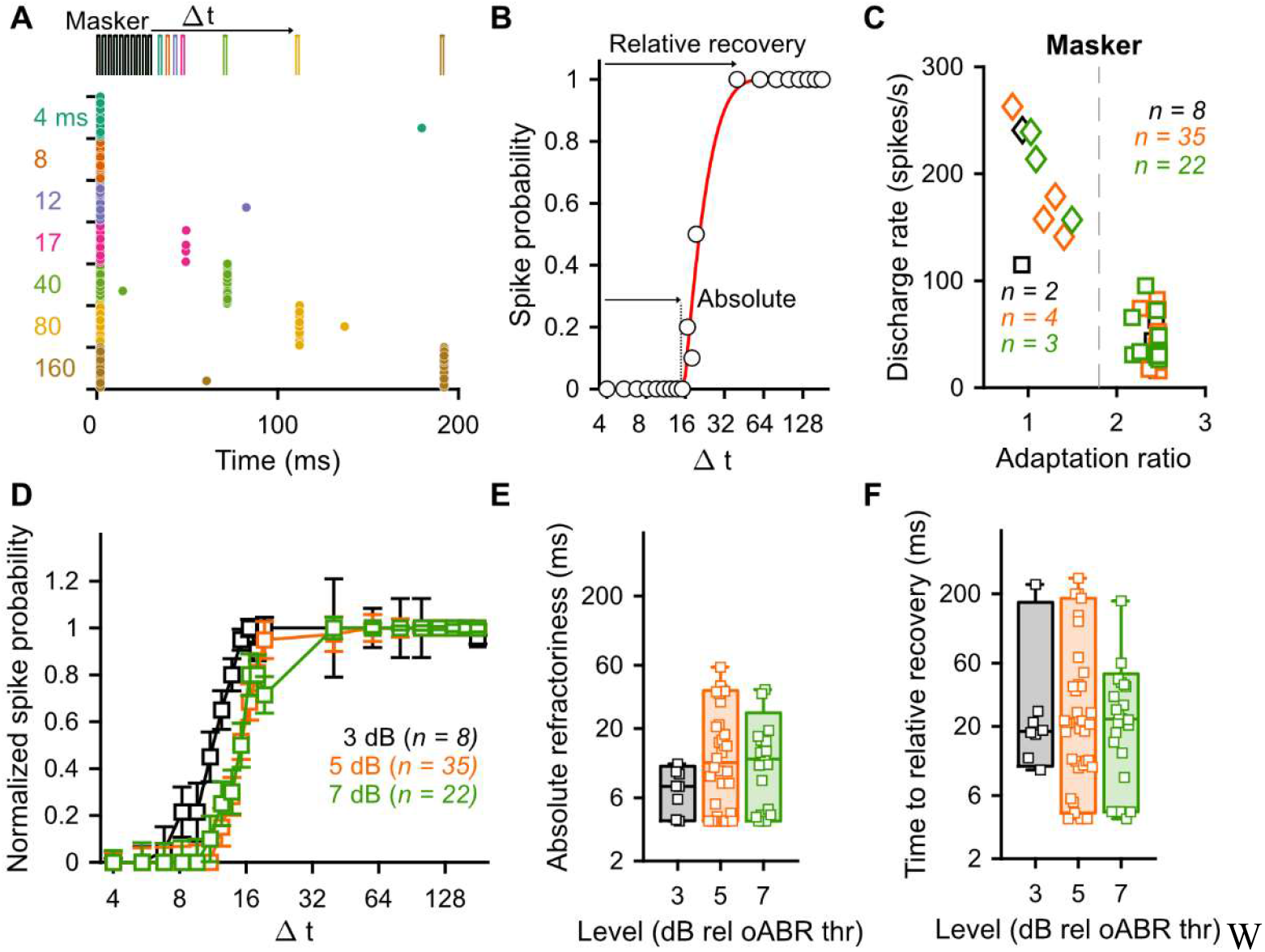
Recovery from optogenetically induced refractoriness is slow and independent of the stimulation level. SGN firing was fully adapted using a masker (10 light pulses, 316 Hz, 1.6 ms duration) and the recovery of firing was assessed at different time intervals (Δ t) ranging between 4 and 200 ms (20 trials, different laser levels ranging between 3 and 6 dB_rel to oABR threshold_, bin width = 1 dB). **A**. Representative raster plot of a SGN undergoing the forward masking protocol at 4.67 dB_rel oABR thr_. A color code is used to represent the different time intervals. **B**. Recovery curve (i.e. spike probability curve as a function of the time interval) of the unit presented in A. The recovery curve was fitted (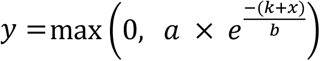 to define the absolute refractoriness (i.e. time required to recover any firing, - k) and time to relative recovery (i.e. time required to recover 95 % of the spike probability, - k + (3 x b)). **C**. The discharge rate as a function of the adaptation ratio allowed to cluster SGNs as non-adapted (i.e. adaptation ratio ≤ 1.8 and discharge rate ≥ 100 spikes/s) and fully adapted. A color code is used for the different levels (D). **D**. The average (± 95% confidence interval) recovery curve measured from fully adapted SGNs at 3 (black, *n* = 8), 5 (orange, *n* = 35), 7 (green, *n* = 22) dB_relative to oABR threshold_ (bin width = 2 dB). **E-F**. Quantification of the time to absolute (E) and relative (F) recovery as a function of the light pulse level for the fully adapted SGNs. The same color code is used as in C. Box plots show minimum, 25th percentile, median, 75th percentile, and maximum with individual data point overlaid. No significant difference was observed between the different tested levels (Jarque-Bera test followed by a Kruskal-Wallis test).

The presence of an optogenetically induced refractoriness, which requires tens to hundreds of milliseconds to recover, highlights the importance of empirically determining the appropriate stimulation parameters for applications to circuit analysis and functional restoration. For example, this information is paramount for studies of the integration of the SGNs firing in complex auditory brainstem neural circuits as well as for designing sound coding strategies for future optical cochlear implants.

### Semi-stochastic stimulation for rapid estimation of the transfer function from light to firing

Empirical estimation of the range of optogenetic stimulus properties appropriate for neural encoding is a tedious task especially if all relevant properties have to be tested for in a deterministic manner which often results in missing data limiting the statistical power of the analysis. To overcome this limitation and rapidly measure the transfer function from light to SGNs firing, we designed a semi-stochastic stimulus (**figure 5.A**) for pairwise testing of 35 combinations of repetition rates (10, 56, 100, 179 and 313 Hz) and pulse durations (0.2, 0.4, 0.6, 0.8, 1.2, 1.6, 2.4 ms). Each combination was randomly tested 10 times per stimulus iteration and the presentation order was randomized for each of the 20 iterations, resulting in 200 presentations per combination. Each iteration started with 10 light pulses of 2.4 ms presented at 100 Hz to adapt the firing and bring the opsin into a desensitized state and finished with 200 ms of the dark. Additionally, a forward masking protocol testing for the intervals of 15 and 80 ms was included. The total duration of the acquisition was 210 s. Recordings were made at maximum radiant fluxes (35 ± 0.35 mW). As with deterministic stimuli (**figure 1**), the optogenetic stimulus was compared to an equivalent acoustic one, recorded from non-AAV-injected mice, composed of acoustical clicks instead of light pulse and presented at various sound levels (60, 70 and 80 dB SPL_PE_).

**Figure 5:**
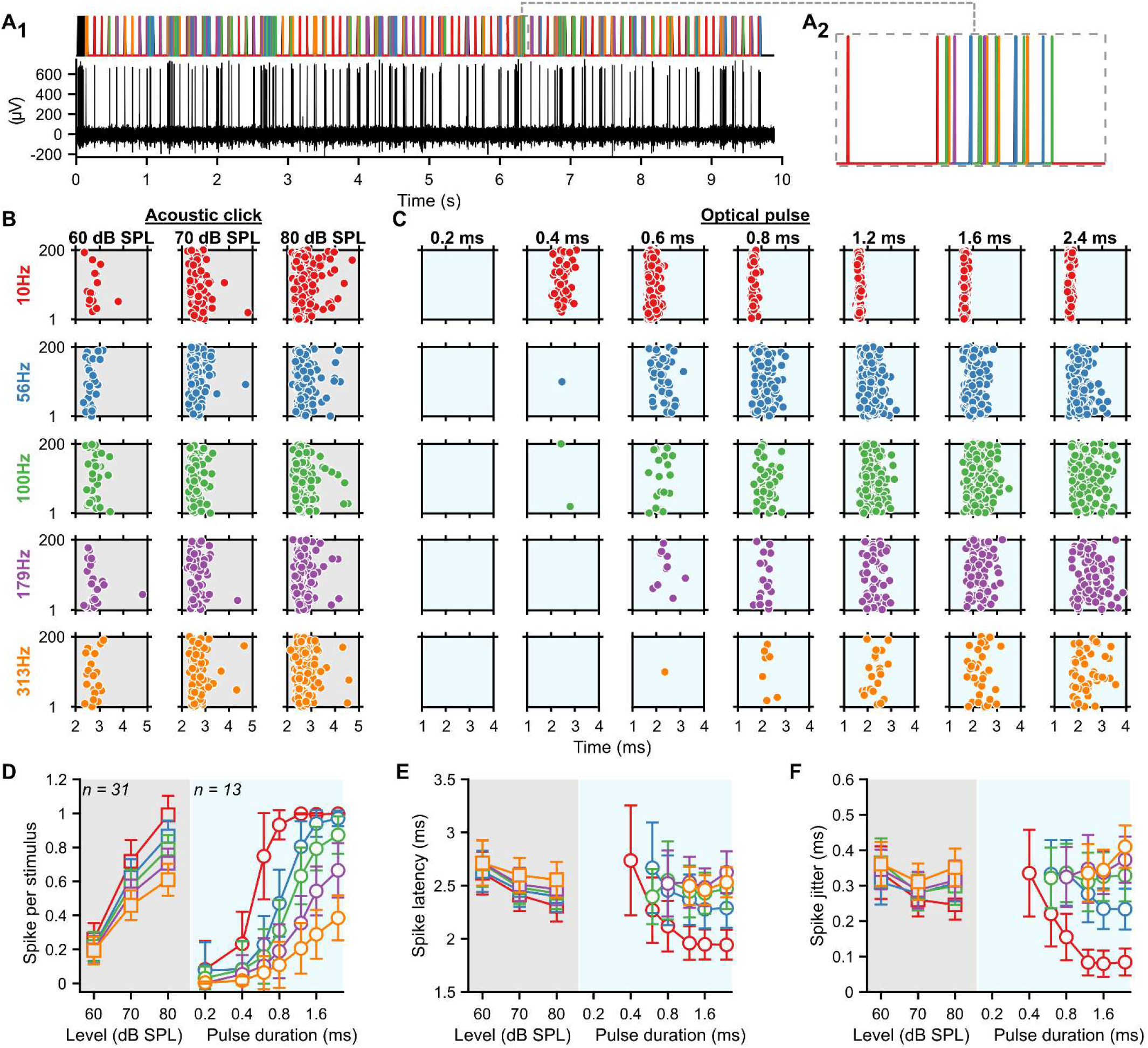
Transfer function from light to SGNs firing can be measured in minutes by a stochastic stimulus. An optical semi-stochastic stimulus (radiant flux = 44 mW) was constructed from 20 presentation of random permutation of repetition rates (10, 56, 100, 179, 313 Hz and color-coded in the figure) and light pulse durations (0.2, 0.4, 0.6, 0.8, 1.2, 1.6, 2.4 ms). The stochastic stimulus was preceded by a masker (10 light pulses of 0.8 ms) and followed by 200 ms of dark per trial. It was compared to an acoustic stimulus constructed from 20 presentations of acoustic click trains containing random permutations of repetition rates (10, 56, 100, 179, 313 Hz) and presented at 60, 70 and 80 dB SPL_PE_. Additionally, a forward masking protocol was included in the optogenetic stimulus with time intervals of 15 and 80ms. **A**. Single iteration of the optical stochastic stimulus (A_1_, top and magnification of the stimulus, A_2_) and the response of representative SGNs (A_1_, bottom). **B-C**. Raster plots from representative acoustic (B, grey background) and optically (C, blue background) evoked (o-)SGNs. **D-F**. Quantification of the number of spikes per stimulus (D), first spike latency (E) and first spike jitter (F) as a function of the sound level for the acoustic modality (grey background, *n* = 31, best frequency within the 16 kHz octave band) and light pulse duration for the optogenetic modality (blue background, *n* = 13). Data are represented as mean and confidence interval (95%).

A total of 13 SGNs were recorded from 2 mice for the optical stimulus and 31 SGNs from 3 mice for the acoustic one. Following reverse correlation and deconvolution (see Methods), raster plots for all tested conditions were built (**figure 5.B-C**) and their spike statistics computed (**figure 5.D-F**). As with the deterministic stimuli, the number of spikes per stimulus increased with the stimulation strength for both modalities (**figure 5.D**). For all light pulse durations or sound intensities, the highest number of spikes per stimulus, shortest latency and highest synchronization were achieved with 10 Hz stimulation (**figure 5.D**). For the optogenetic modality, higher repetition rates required longer light pulses to evoke the same number of spikes per stimulus and significantly influenced the number of spikes evoked per light pulse and their timing (**figure S4** for pairwise comparisons of the repetition rates as a function of the pulse duration). Interestingly, reaching the same number of spikes per stimulus with rates > 10 Hz was always associated with longer latency and higher jitter (**figure 5.E-F**). Thus the stochastic stimulus also allows evoking spike trains with the same discharge rate but different spike timing in SGNs.

### Heterogeneous optogenetic activation of the SGNs

We report heterogeneous activation of the SGNs whereby the number of spikes evoked per light pulse and their temporal precision ranged over 1 order of magnitude (**figure 2**), activation thresholds over 7 dB, dynamic range over 9 dB (**figure 3**) and recovery time from refractoriness over 200 ms (**figure 4**). To investigate the mechanisms underlying this heterogeneity, we capitalized on 11 SGNs that were recorded from the same animal, one after the other, in the same condition of optical stimulation and recordings. All neurons were recorded in response to the semi-stochastic stimulus presented above and a protocol testing forward masking for intervals of 15 and 80 ms. A principal component analysis of the number of spikes per light pulse for the 35 tested conditions highlighted the presence of 3 clusters (**figure 6.A-B**, 91.4% of the variance explained by principal components 1 and 2). Encoding of the light pulse differed between the three clusters with SGNs from cluster 3 being the most efficient to encode the light pulses and cluster 1 the worst (**figure 6.C-D**). The light pulse threshold (i.e. the shortest light pulse required to evoke 0.1 spikes per light pulse) was significantly lower for clusters 2 and 3 compared to cluster 1 and linearly increased with the repetition rate (**figure 6.E**, *p*-values ranging between 0.0056 and 0.0278, Kruskal-Wallis test followed by a Tukey’s Honestly Significant Difference Procedure). The linear regression slope was significantly lower for cluster 3 compared to cluster 1 (**figure 6F**, *p*-value = 0.0138), likely reflecting differences between clusters in the light dependence of the photocurrent amplitude on the repetition rate.

**Figure 6:**
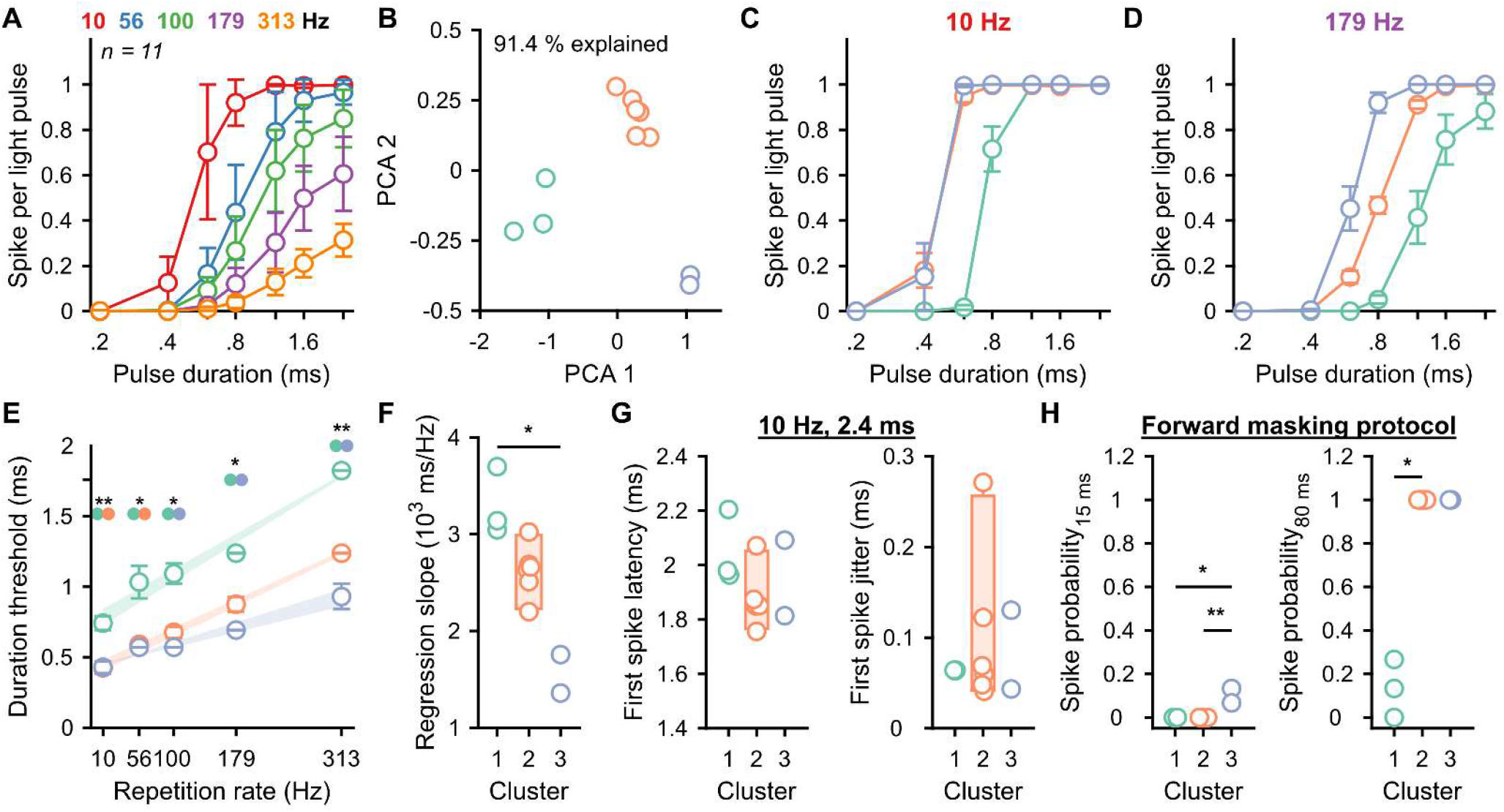
Heterogeneous responses among SGNs within a single animal. **A**. Quantification of the number of spikes per light pulse as a function of the repetition rate and light pulse duration from 11 SGNs measured from the same animal. Data are represented as mean and confidence interval (95%). **B**. The numbers of spike per light pulse measured at all conditions of repetition rates and light pulse durations were used to perform a principal component analysis (PCA, first 2 components explain 91.4 % of the variance) which highlight the presence of 3 SGNs clusters (cluster 1, green, *n* = 3; cluster 2, orange, *n* = 6; cluster 3, purple, *n* = 2). **C-D**. Quantification of the number of spikes per light pulse for the 3 SGN clusters as a function of the light pulse duration at 10 (C) and 179 Hz (D). Data are represented as the mean and standard error of the mean. **E**_**1**_. Quantification of the light pulse duration threshold as a function of the repetition rate per cluster. The average linear regression (and standard error of the mean) between threshold and repetition rate is represented by the light fill. The threshold difference between clusters was tested using a Kruskal-Wallis test followed by a Tukey’s Honestly Significant Difference Procedure (*, p-value ≤ 0.05). **E**_**2**_. Quantification of the light pulse threshold dependency to the repetition rate. Box plots show minimum, 25th percentile, median, 75th percentile, and maximum with individual data point overlaid. The difference between clusters was tested using a Kruskal-Wallis test followed by a Tukey’s Honestly Significant Difference Procedure (*, p-value ≤ 0.05; **, p-value ≤ 0.01). **F**. Quantification of the first spike latency (left) and first spike jitter (right) measured in response to the condition repetition rate = 10 Hz and light pulse duration = 2.4 ms per cluster. No difference was measured between clusters (Kruskal-Wallis test). **G**. Quantification of the spike probability recovery (forward masking protocol as described in figure 3) at 15 ms (left) and 80 ms (right) per cluster.

Surprisingly, the latency and jitter of the first spike evoked by light pulses of 2.4 ms at 10 Hz were similar across clusters which seems to contrast with previous results demonstrating shorter spike latencies with a greater current amplitude injected into the postsynaptic bouton of SGNs (Rutherford et al., 2012). Finally, SGNs from cluster 3 were the fastest to recover from refractoriness (spike probability at 15 ms, 0 ± 0, 0 ± 0 and 0.1 ± 0.42 spike for cluster 1, 2 and 3 respectively, average ± 95% confidence interval, *p*-values ranging between 0.0073 and 0.0193), followed by cluster 2 and 1 (**figure 6.H**, spike probability at 80 ms, 0.13 ± 0.33, 1 ± 0 and 1 ± 0 spike for cluster 1, 2 and 3 respectively, *p*-values ranging between 0.008 and 0.054). Together these results demonstrate that SGNs encoding light stimuli the best also are the ones recovering the fastest from the refractoriness. Hence, we postulate that those neurons are the ones with the largest photocurrents (see Discussion).

### Enhanced representation of the optogenetic stimulation in AVCN neurons

Above, we reported that the semi-stochastic stimulus allowed generation, in the same neurons, of spike trains with different statistics in terms of the number of spikes and spike timing (latency and jitter, **figure 5**). Here, we investigated if this property could be used to study the information transfer from the SGN to AVCN neurons. First, we observed that the success of given optogenetic stimulus to trigger a spike depended on the success/failure of the preceding light pulse for all tested repetition rates. If the preceding light pulse had a sub-threshold duration, the likelihood of the neuron to encode the following light pulse was high. In contrast, if the previous light pulse had a duration above the threshold, the likelihood to fire upon the following pulse was low. This explains the sawtooth pattern of the response in **figure 7.A-B**. Therefore, spike statistics were computed separately for both modalities as a function of the dark/silence time (i.e. inter stimulus interval). This resulted in 245 illumination conditions and each of them was tested 15.71 ± 0.42 times for each SGN (average ± 95% confidence interval), whereas we did not change the number of acoustical conditions.

**Figure 7:**
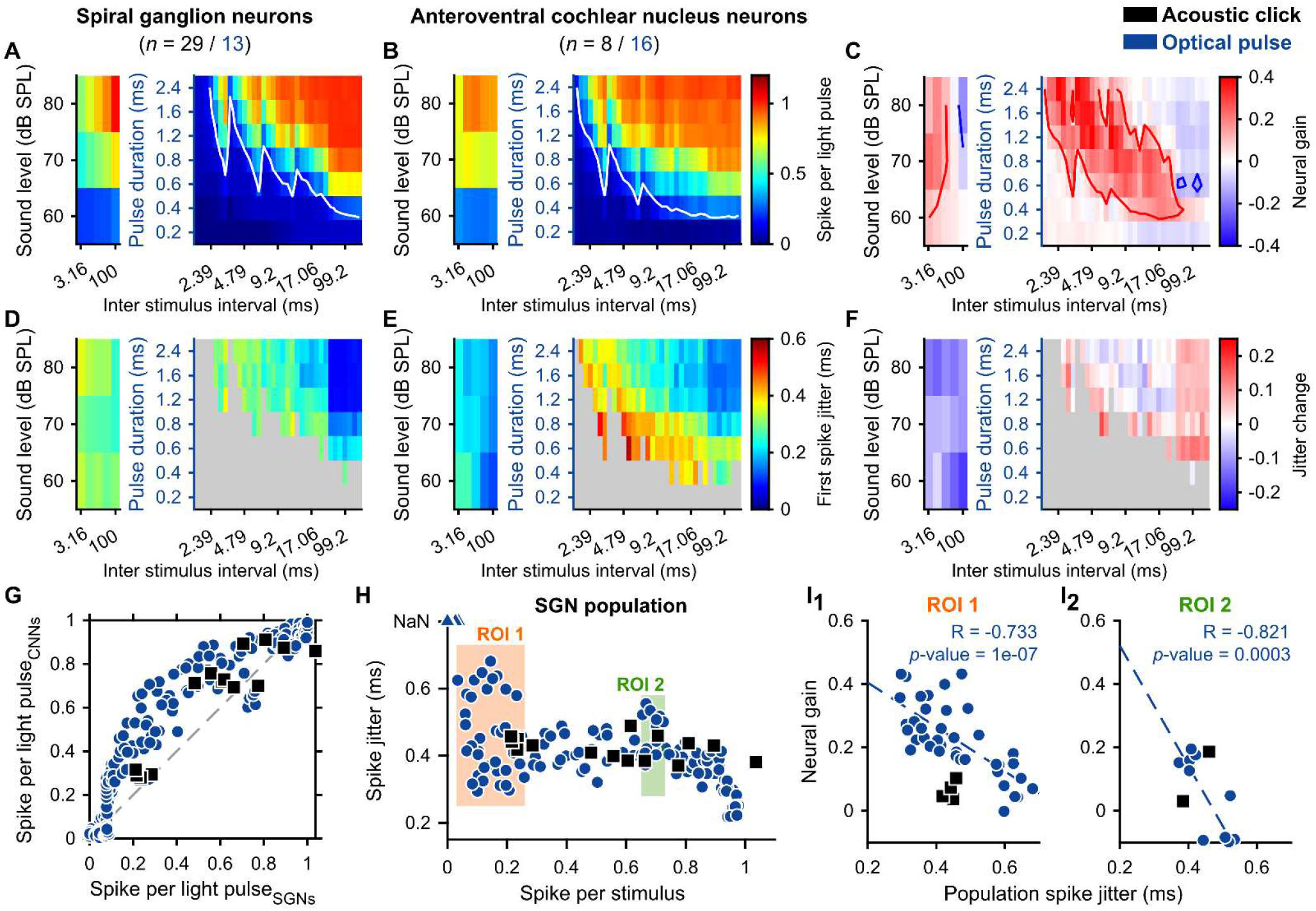
The neural gain of the AVCN neurons depends on the level of synchronization between SGNs. The initial 35 conditions of repetition rates and light pulse durations were divided into 235 conditions of inter-stimulus intervals and pulse durations. Note that all conditions were tested randomly and in parallel for all neurons. **A-B, D-E**. Quantification of the number of spikes per stimulus (A-B) and first spike jitter (D-E) for the different tested conditions for SGNs (A, D, acoustic click, black, *n* = 29; light pulse, blue, *n* = 13) and AVCN neurons (B, E, acoustic click, *n* = 8; light pulse, *n* = 16). In A-B, the white line delimits the threshold defined as 0.2 spike/stimulus. **C-F**. Quantification of the neural gain (C) and jitter change (F) between SGNs and AVCN neurons. In C, the red line delimits the area where the neural gain is ≥ 0.1 and the blue line where the neural gain is ≤ 0.1. **G**. Average number of spikes per light pulse of the AVCN neurons as a function of the number of spikes per light pulse of the SGNs. A blue circle (optogenetic) or a black square (acoustic) corresponds to one of the stimulation conditions tested for all neurons. **H**. Average first spike jitter as a function of the number of spikes per light pulse computed across all SGNs per stimulation condition. Two regions of interest (ROI) were defined as the region where the numbers of spikes per light pulse were in the same range and the variation of the first spike jitter was large (ROI 1, 0 ≤ number of spikes per stimulus ≤ 0.26; ROI 2, 0.65 ≤ number of spike per stimulus ≤ 0.73). **I**. Average neural gain as a function of the first spike jitter across SGN in ROI 1 (I_1_) and ROI 2 (I_2_). Correlation coefficients (R and its significance) were measured and displayed as a straight line if *p*-value ≤ 0.05.

To simplify the visualization, the average number of spikes per stimulus and the first spike latency jitter were color-coded for SGNs (**figure 7.A,D**, optical: *n* = 13 from 2 mice; acoustical: *n* = 26 from 4 mice, CF within the 16 and 32 kHz octave band) and AVCN neurons (**figure 7.B,E**, optical: *n* = 16 from 5 mice; acoustic: *n* = 8 from 3 mice, CF within the 16 and 32 kHz octave band). As previously described the best encoding was achieved at the longest inter-stimulus interval and stimulation strength. The change in the response between SGNs and AVCNs was highlighted by computing the difference between the two (**figure 7.C,F**). For 38.37% of the optical and 46.67% of the acoustic conditions, the number of spikes per stimulus was increased by at least 0.1 spike/stimulus (i.e. positive neural gain) in the AVCN neurons compared to the SGNs. In contrast, the response decreased by 0.1 in 1.22% of the optical and 6.67% of the acoustic conditions (**figure 7.C,G**). The spike jitter decreased for all acoustic tested conditions, and stayed similar for most of the optogenetic ones (**figure 7.F**).

Presynaptic terminals of the SGNs converge on the AVCN somas where their glutamatergic inputs trigger post-synaptic action potentials (Cao and Oertel, 2010; Strenzke et al., 2009). To interpret the neural gain associated with the different spike statistics of the SGNs, it is required to interpret their firing from the perspective of the postsynaptic neurons receiving their convergent inputs. To do so, population responses were computed across recorded SGNs (see Methods, **figure 7.H**). For both modalities, the jitter across SGNs tended to increase with the number of spikes per light pulse in the AVCN neuron, but was constant between 0.3 and 0.9 spikes per light pulse. Based on the optogenetic response, two regions of interest (ROI) were defined where the number of spikes was constant but the jitter varied across SGNs (**figure 7.H**). For both ROIs, in the optogenetic modality, the neural gain was negatively correlated with the jitter (ROI 1: coefficient of correlation, R = -0.733, *p*-value = 1 × 10^−7^; ROI 2: R = -0.821, p-value = 3 × 10^−4^), suggesting that synchronous SGNs inputs on AVCN neurons are required to increase their rate representation of the stimulus (**figure 7.I**).

## Discussion

Here, we parametrized optogenetic intensity coding in SGN firing mediated by the ultra-fast, trafficking-optimized Chronos and the recoding of information by AVCN neurons. We demonstrate graded optogenetic SGN activation with varying pulse widths like SGN responses to acoustic clicks with different sound pressure levels. We find that the timing of optogenetically-evoked spikes varied little across iterations for a given SGN, but across the SGN population is similar to what is observed acoustically. We identified an effective light pulse width of 1.6 ms to optimally encode the light intensity (radiant flux) achieving the highest maximal discharge rate and the widest dynamic range. Upon high rates of stimulation, we observed robust spike rate adaptation that often eventually resulted in spike failure. We attribute this to a depolarization block which required a few tens of milliseconds to recover independently of the tested light intensities. Finally, we implemented rapid measurements (minutes) of the transfer function from light to SGN firing rate using a semi-stochastic stimulus. Spike statistics in the SGN population (i.e. number of spikes and spikes timing) generated by this stimulus also enabled us to study the recoding of the spike trains by the AVCN neurons. We observed that some synchronization between SGNs is required to enhance the rate response of AVCN neurons via integrating their excitatory inputs.

### Achieving gradual Chronos-mediated activation of the SGNs with the pulse width

A dependence of photocurrent amplitude on the photon flux has already been shown in early studies of algal rhodopsins (Harz and Hegemann, 1991) and therefore manipulating the light pulse amplitude or its width enables one to control of the number of action potentials mediated by those photocurrents (e.g. (Berndt et al., 2011; Boyden et al., 2005; Gunaydin et al., 2010; Klapoetke et al., 2014; Mager et al., 2018; Zhang et al., 2008)). The application of this concept to gradually activate neural network activity, to our knowledge, has not been widely employed. In combination with an electrophysiological or optical read-out of the neuronal response, it would enable measuring the transfer function between one element of a given network to another. Moreover, it is also of prime importance to establish an appropriate stimulation parameter space for sensory coding strategies to draw from when implement future optogenetic prosthetics, such as the optical cochlear implant (Dieter et al., 2020; Wolf et al., 2022). Both applications require empirical measurements of the optogenetically evoked spiking which depends on the opsin biophysics (Klapoetke et al., 2014; Mager et al., 2018; Mattis et al., 2011), opsin expression (Meng et al., 2019) and neuron excitability (Herman et al., 2014; Mager et al., 2018). Using the auditory system as a model circuit with exquisite temporal fidelity, here, we confirmed with a single light pulse that gradual activation of SGNs expressing Chronos-ES/TS can be achieved by individual light pulses up to 1.15 spikes/pulse using long light pulse durations (i.e. 3.2 ms, **figure 1.E**). Interestingly, for pulse trains (100 Hz), the number of spike/pulse peaked at 0.74 for pulses of 1.6 ms and decreased for shorter or longer light pulse durations (**figure 1.J**). It is also for the longest light pulse that the spike adaptation was the strongest, which contrasts with the fact that such adaptation was not observed for acoustically-driven SGNs (**figure S2.B**). This suggests that the dark time between two consecutive pulses might have been too short to enable recovery of Chronos-mediated photocurrent.

### Recovery from light-induced refractoriness

Sustained firing could not be recorded from any of the SGNs when stimulated by trains of light pulses at high repetition rates (i.e. 316 and 1000 Hz, **figure S2.C,E**). This observation was unexpected given the fast closing kinetics reported for Chronos (Keppeler et al., 2018; Klapoetke et al., 2014). However, such rapid spike rate adaptation is consistent with our previous recordings of SGN firing mediated by fast opsins (Chronos-ES/TS (Keppeler et al., 2018) and vf-Chrimson (Bali et al., 2021)). Current-clamp recordings of SGNs have revealed that most of the SGNs are characterized by a rapidly- or intermediate-adapting firing in response to current injections (Markowitz and Kalluri, 2020; Mo and Davis, 1997). Other studies have shown that neurons rapidly adapting to a current injection are likely to have an optogenetically induced depolarization block using long light pulse durations or high stimulation rates (Herman et al., 2014; Mattis et al., 2011). We investigated the time needed to recover from the depolarization block using a so-called forward masking protocol. The masker could induce complete adaptation in approximately 85% of the recorded SGNs, which is similar to the number of rapidly adapting SGNs reported by patch-clamp studies on cultured neonatal SGNs (Mo and Davis, 1997). The time of recovery from the full adaptation of those neurons amounted to ∼30 ms which is consistent with the time their voltage-gated sodium channels require to recover from inactivation (Lin, 1997). The best strategy to reach high firing rates of fast-adapting neurons has yet to be elaborated and likely should aim to eliminate the plateau potential leading to the sodium channel inactivation (as previously shown with ChETA in fast-spiking neurons, (Gunaydin et al., 2010). Possibilities include the activation of fast-gating hyperpolarizing opsins such as anion permeating ChRs (Govorunova et al., 2015) right after optogenetically triggering a spike.

### Optogenetic encoding of intensity information

Encoding the intensity of a stimulus is critical in the context of sensory restoration. Optically it corresponds to the number of photons reaching ChRs which can be controlled by the pulse width and/or the light radiant flux. We previously examined the radiant flux range associated with a change in the firing rate of the SGNs expressing the red-shifted f- and vf-Chrimson (Bali et al., 2021). We reported that the dynamic range (in dB) for single neurons spanned between 0.67 and 3.77, seemed to be opsin-dependent but was less than measured acoustically (∼ 25 dB (SPL)). Here we investigated the co-variation of the radiant-flux and the pulse width to investigate how they jointly impact the intensity encoding. Using 1.6 ms light pulse we achieved an optimal intensity encoding defined as the lowest threshold (1.50 dB_rel to oABR threshold_) and the widest dynamic range (2.26 dB) and driven rate (84.45 spikes/s, **figure 3**). This duration is consistent with ChR2’s lifetime of the open state (∼1.5 ms), during which the opsin is integrating photons (Ritter et al., 2008; Saran et al., 2018). A longer light pulse could accommodate more than one ChR opening, likely increasing the variance and adaptation of the photocurrents.

### Optogenetic encoding of temporal information

Reliable optogenetic encoding of the spike timing is critical to resolving neural circuits for which time matters (Cariani and Baker, 2022), such as the auditory system where spike timing is critical to localize horizontally low-frequency sounds, discriminate the pitch or detect sound in noise (Grothe et al., 2010; Huet et al., 2018). Our current and previous works have shown a significantly higher temporal precision of firing for optogenetic than for acoustic stimulation when evaluated at a single neuron level and which requires careful interpretation (**figure 1.G,L**, (Bali et al., 2021)). Indeed, this high temporal precision is measured for a given neuron in response to multiple iterations of the same stimulus, which can be interpreted as a highly reliable encoding of the light pulses. Importantly, here, we report that the latency at which neurons are firing is heterogeneous (**figure 1.F,K, figure 2.B, figure 6.G**) and therefore the excitatory inputs from a population of SGNs into postsynaptic neurons might be less precisely timed than initially estimated. As an attempt to address this question, we computed the precision of the spikes within one population of SGNs built from all the recordings we performed and compared it to acoustically evoked SGNs (**figure 2.D-E**). This analysis revealed comparable spike time precision between the two modalities, which suggests that postsynaptic neurons might be excited in a near-physiological manner optogenetically.

### Transfer function from light to SGN firing

Measuring empirically the illumination parameters required for a given application of optogenetics in neural network analysis or sensory restoration is a prerequisite, as the variability of neuronal excitability, opsin biophysical properties and expression levels can challenge predictions (Gunaydin et al., 2010; Herman et al., 2014; Jun and Cardin, 2020; Mager et al., 2018; Mattis et al., 2011). Estimating those parameters using deterministic stimuli is nearly impossible given the limited stability and time scope of recordings such as of SGN firing (e.g. in vivo juxtacellular recordings of the SGN axons). Thus, only enabling so far sparse estimation of the transfer function from light to SGN firing (Bali et al., 2021; Hernandez et al., 2014; Keppeler et al., 2018; Mager et al., 2018; Wrobel et al., 2018). Using a semi-stochastic stimulus containing parameters of interest allowed us to measure in minutes (∼ 200 s) 35 conditions of repetition rates and light pulse duration (or, 245 conditions of dark interval and light pulse duration) to estimate the transfer function (**figure 5**). By doing so, we replicated the results obtained from the deterministic approach we previously used (**figure 1**) and measured intermediate values. The number of conditions measured pairwise per neuron enabled us for the first time to identify 3 distinct clusters of SGNs which differed in their encoding of the light pulses and their recovery of the depolarization block (**figure 6**). The neurons requiring less light to fire were the same ones recovering the fastest from the depolarization block potentially suggesting a difference in expression level or neuron excitability. Future studies should address the underlying mechanisms by, for example, labeling SGNs to discriminate between molecular or morphological differences between SGNs (Liberman, 1982; Petitpré et al., 2018; Shrestha et al., 2018; Sun et al., 2018), the difference of the expression levels (Meng et al., 2019), or the difference of distance to the light emitter.

### Anteroventral cochlear nucleus neurons integration

Axons of the SGNs are converging on the somas of different cell types where they provide strong excitatory inputs (Cao and Oertel, 2010; Liberman, 1991; Strenzke et al., 2009; Tsuji and Liberman, 1997). Within the AVCN, the bushy cells increase the probability and precision of their cycle-to-cycle response to low-frequency sounds (Joris et al., 1994; Wei et al., 2017) but the underlying mechanisms of this enhancement are still under debate (Ashida et al., 2019; Joris and Smith, 2008). We took advantage of the heterogeneous SGNs firing evoked by all conditions of the semi-stochastic optogenetic stimulation aiming to identify the transfer function from SGN to AVCN neuron firing. We observed acoustically and optogenetically an increase in the firing of the AVCN neurons compared to the SGNs in some of the tested conditions (**figure 7.A-C**) which was associated with no or little change in the spike precision. Optogenetically, the observed neural gain was correlated to the level of synchronization across SGNs suggesting that coincident inputs were required to increase the discharge rate of the AVCN neurons (**figure 7.G-I**), while no correlations were observed acoustically. The precise mechanisms underlying this result are unclear and future studies will require to group AVCN neurons according to their cell types (e.g. spherical bushy cell, globular bushy cell, stellate cells, …) thereby taking into account their known morphology and physiology when interpreting the data. Nonetheless, this result supports the concept that employing additional stimulation modalities such as optogenetics will help to reveal unknown properties of the elements constituting neural circuits and contribute to their understanding.

### Lessons learned for optogenetic information coding

The results inform efforts for improved functional restoration in diseased sensory and motor systems by optogenetic neural stimulation. As Chronos is likely the fastest optogenetic actuator and SGNs serve as targets exemplifying very high temporal fidelity of coding, the strong spike rate adaptation observed indicates an upper bound of at stimulus rates of ∼200 Hz (our data and (Keppeler et al., 2018)), at least for the photocurrents amenable from algal ChRs and for continuous stimulus encoding. Moreover, high intensity light pulses with graded effective durations in range of 400 μs to 2 ms seem appropriate as the “fundamental” stimulus. Considering laser diode operation being most efficient for μs short and tens of mA strong driver currents, this suggest to synthesize such effective fundamental stimuli from μs pulselets. The *in vivo* data presented here indicate that SGNs readily integrate the photocurrents elicited by such pulselets, which seems in line with previous patch-clamp studies on spike generation (Markowitz and Kalluri, 2020; Mo and Davis, 1997; Rutherford et al., 2012; Smith et al., 2015). It will be of interest for the ultimate design of optogenetic sound coding strategies to carefully parametrize the effects of pulselet duration and duty cycle of such synthetic stimuli in future experimental and theoretical studies. Likewise, challenging patch-clamp recordings from ChR expressing SGNs (e.g. Mager et al., 2018) will be needed to elucidate the refractoriness to optogenetic stimulation. Regardless of the precise underlying mechanisms, this results places an upper limit on the stimulation rate for reliable sound encoding. Finally, the present study reveals diversity of optogenetically driven SGN firing that extends information coded by the population well beyond the coding capacity of the individual SGN. From a theoretical perspective, neuronal functional diversity across a population of neurons is a key factor to widen the amount of encoded information and increase its fidelity (Berry Ii et al., 2019). The presence of functionally different SGNs clusters obtained with administration of a single viral vector-promotor-ChR combination provides a flavor of what could be obtained when targeting different ChRs to molecularly distinct SGN type I subtypes. While preparing a first clinical trial suggests to avoid the potential risk generated by additional complexity, future generations of optogenetic hearing restoration could capitalize on these exciting opportunities.

---

**Figure S1:**
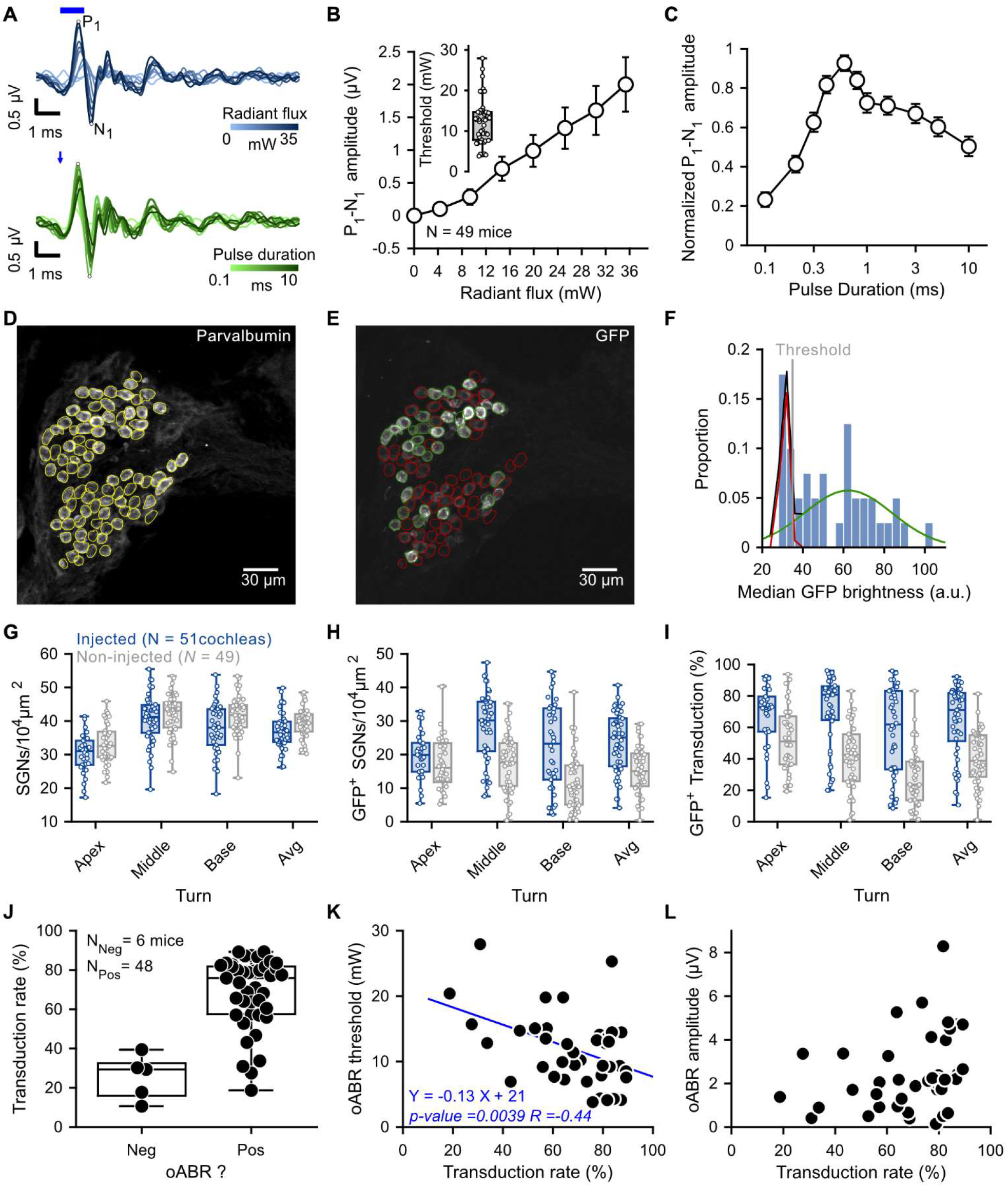
Physiological and histological characterization of the optogenetic modification of murine spiral ganglion neurons using Chronos-ES/TS. **A**. Representative optogenetically evoked auditory brainstem responses (oABR) to various radiant flux (top panel, blue, 1 ms light pulses presented at 10 Hz, the tick blue line represents the light pulse) and light pulse durations (lower panel, green, 35 mW light pulses presented at 10 Hz, the blue arrow represents stimulus onset). The first wave complex (P_1_-N_1_) corresponds to the synchronous activation of the SGNs. **B-C**. Quantification of P_1_-N_1_ amplitude (*N* = 49 mice) as a function of the radiant flux (B, radiant flux binned per 5 mW steps) and pulse duration (**C**.). Average ± 5% confidence interval. Inset in B.: Box-and-whisker plot (minimum, 25^th^, median, 75^th^ percentile, maximum) of oABR threshold. **D-F**. Chronos-ES/TS transduction was semi-automatically evaluated from confocal images of mid-modiolar cryosection stained for Parvalbumin (D) and GFP (E). The method was previously described in (Huet et al., 2021) and consisted to segment individual SGN somas from the Parvalbumin images, measuring the median GFP signal from individual segmented SGNs and defining, using a Gaussian-mixture model, the threshold (vertical grey line) at which the median GFP brightness exceeded (average + 3 standard deviations, green curve) the distribution of the GFP background noise (F, red curve). **G-I**. Quantification of the SGN density (i.e number of parvalbumin-positive SGN somas per cross-sectional area of Rosenthal’s canal, G), the density of GFP positive SGNs (H), and transduction rate (i.e. the ratio between the GFP positive SGN density and the SGN density, I) for all cochlear turns of injected (blue, *N* = 51 cochleae) and contralateral non-injected (grey, *N* = 49) mouse cochleae. Box plots show minimum, 25th percentile, median, 75th percentile, and maximum with individual data point overlaid. **J**. Quantification of the transduction rate as a function of the oABR result. **K-L**. Correlation of the transduction rate with the oABR threshold (K) and oABR N_1_-P_1_ amplitude (L). In case a significant correlation coefficient was measured, a linear model (blue line) was fitted to the data.

**Figure S2:**
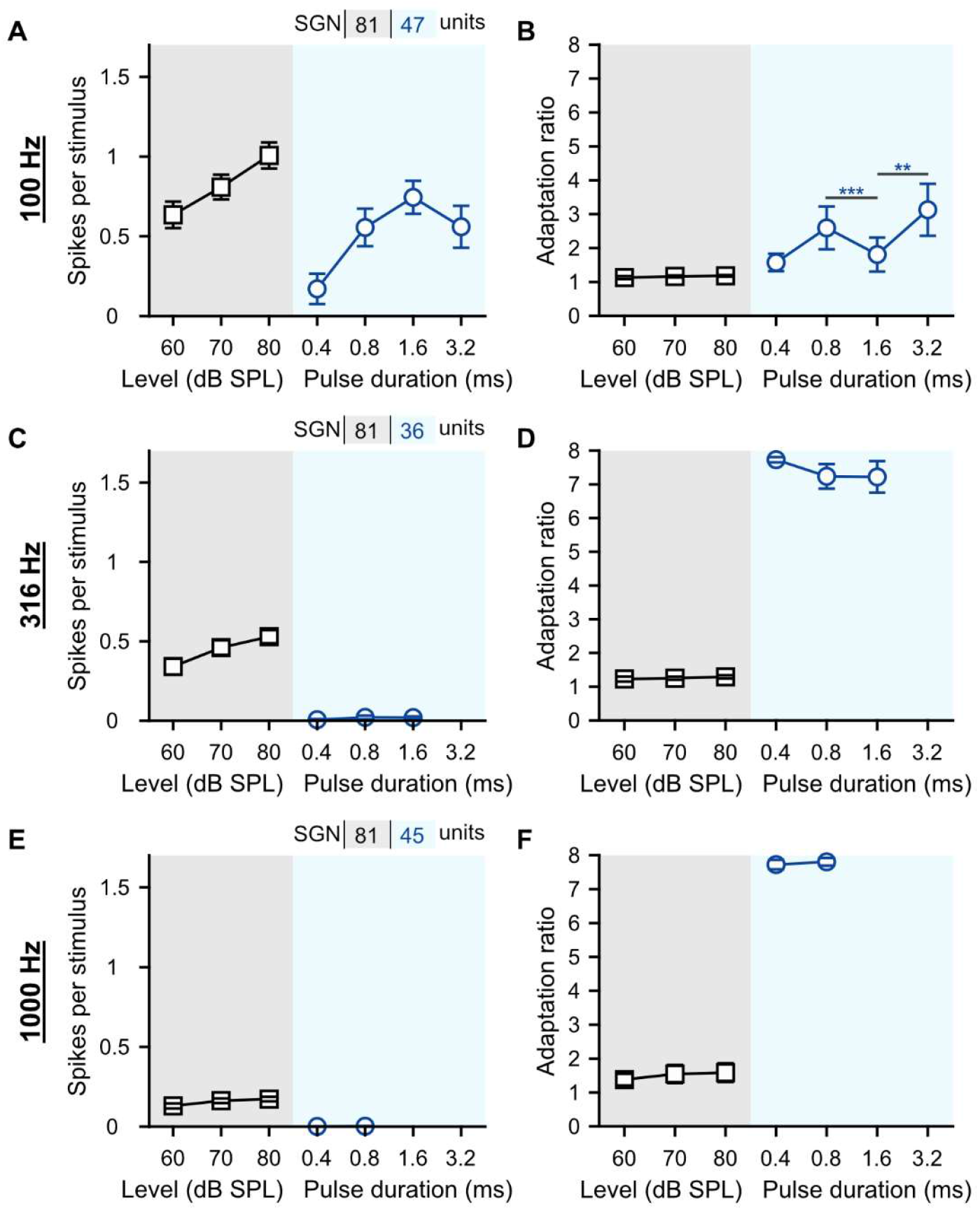
316 and 1000 Hz light pulse trains fully adapted the SGN firing. Quantification of the number of spikes per stimulus (**A, C and E**) and adaptation ratio (i.e. the ratio of the discharge rate during the first 400 ms and the first 50 ms of the stimulation, **B, D and F**) at 100 (**A-B**), 316 (**C-D**) and 1000 Hz (**E-F**). The effect of the acoustic levels and light pulse durations was tested by a Wilcoxon signed rank test on paired samples followed by a Bonferroni correction of the *p*-values (**, *p*-value ≤ 0.01; ***, *p*-value ≤ 0.001).

**Figure S3:**
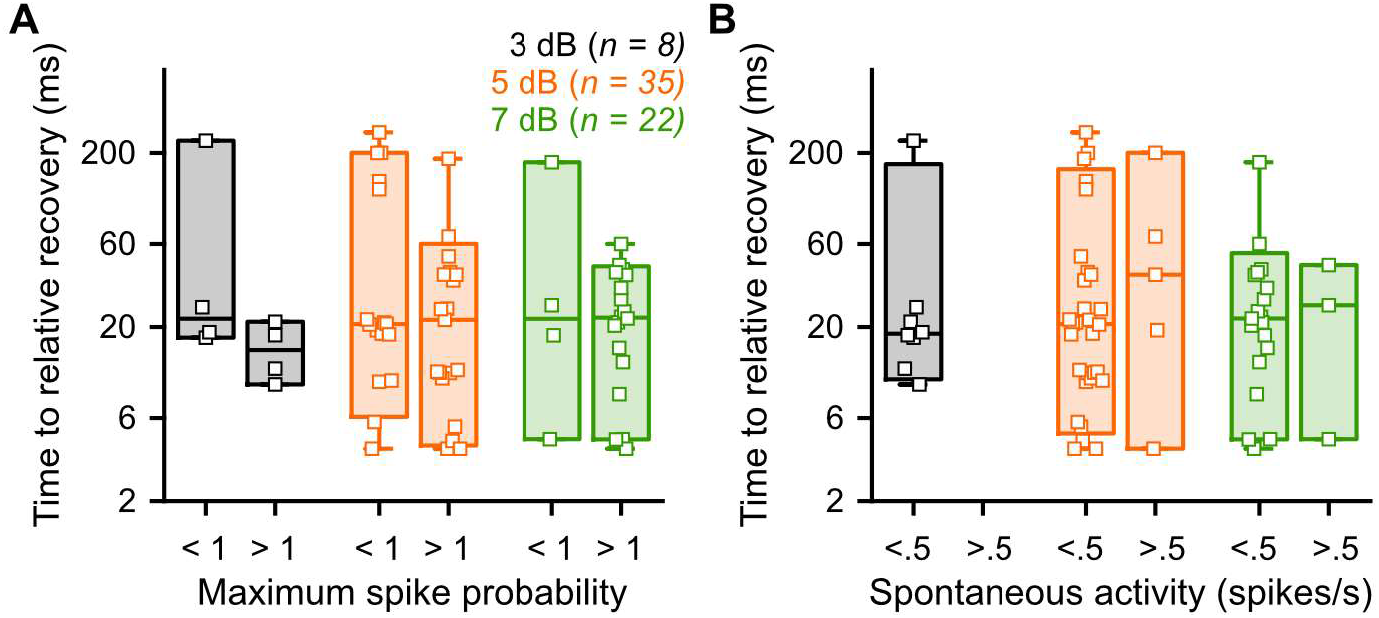
Time to relative recovery is independent of the spontaneous activity (A) and maximum spike probability (B). The spontaneous activity was measured in the last 200 ms following the longest tested interval (Δ t). The maximum spike probability was the spike probability measured in response to the probe eliciting the highest spike probability. Light levels were expressed as dB_relative to oABR threshold_ (bin width = 2 dB) and color-coded.

**Figure S3:**
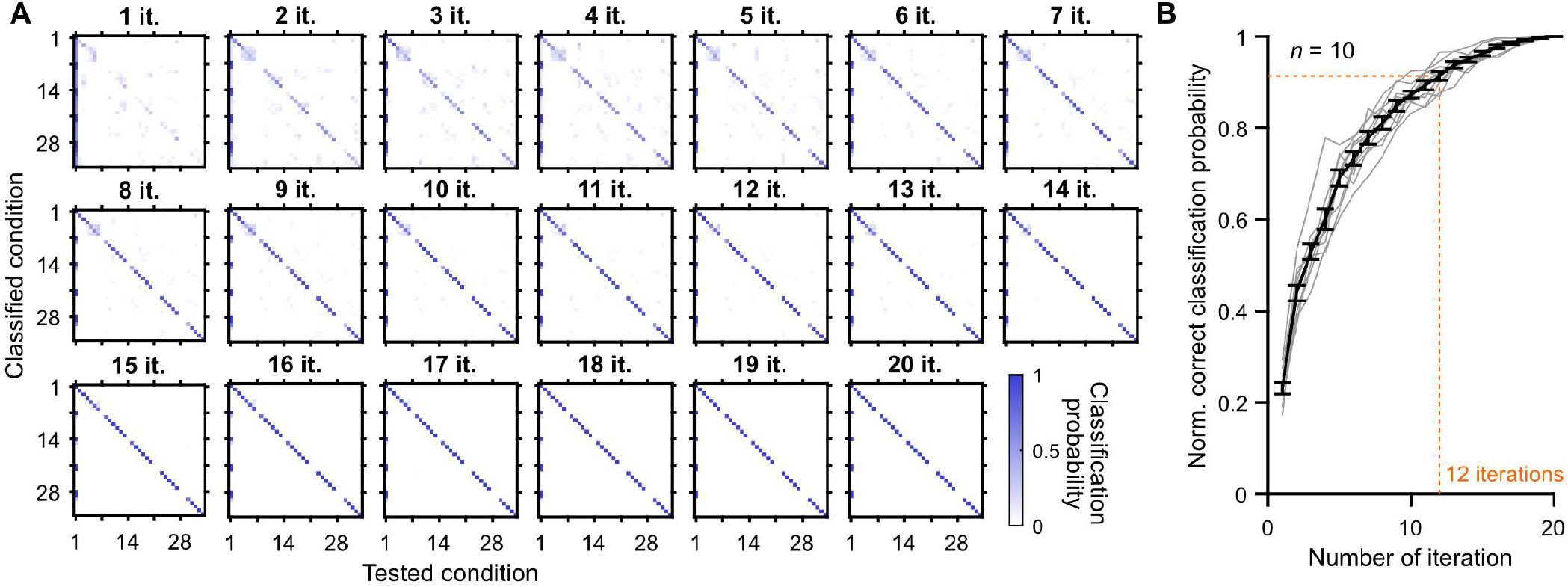
12 iterations are needed per condition for the semi-stochastic stimulus. Subplots are showing the classification probability of 35 tested and classified conditions (7 pulse durations x 5 repetition rates in the semi-stochastic stimulus) for their repetitive presentations (10-200). The plot shows the normalized correct classification probability as a function of the number of presentations. On average 12 iterations are needed to have a correct classification probability of 1.

**Figure S4:**
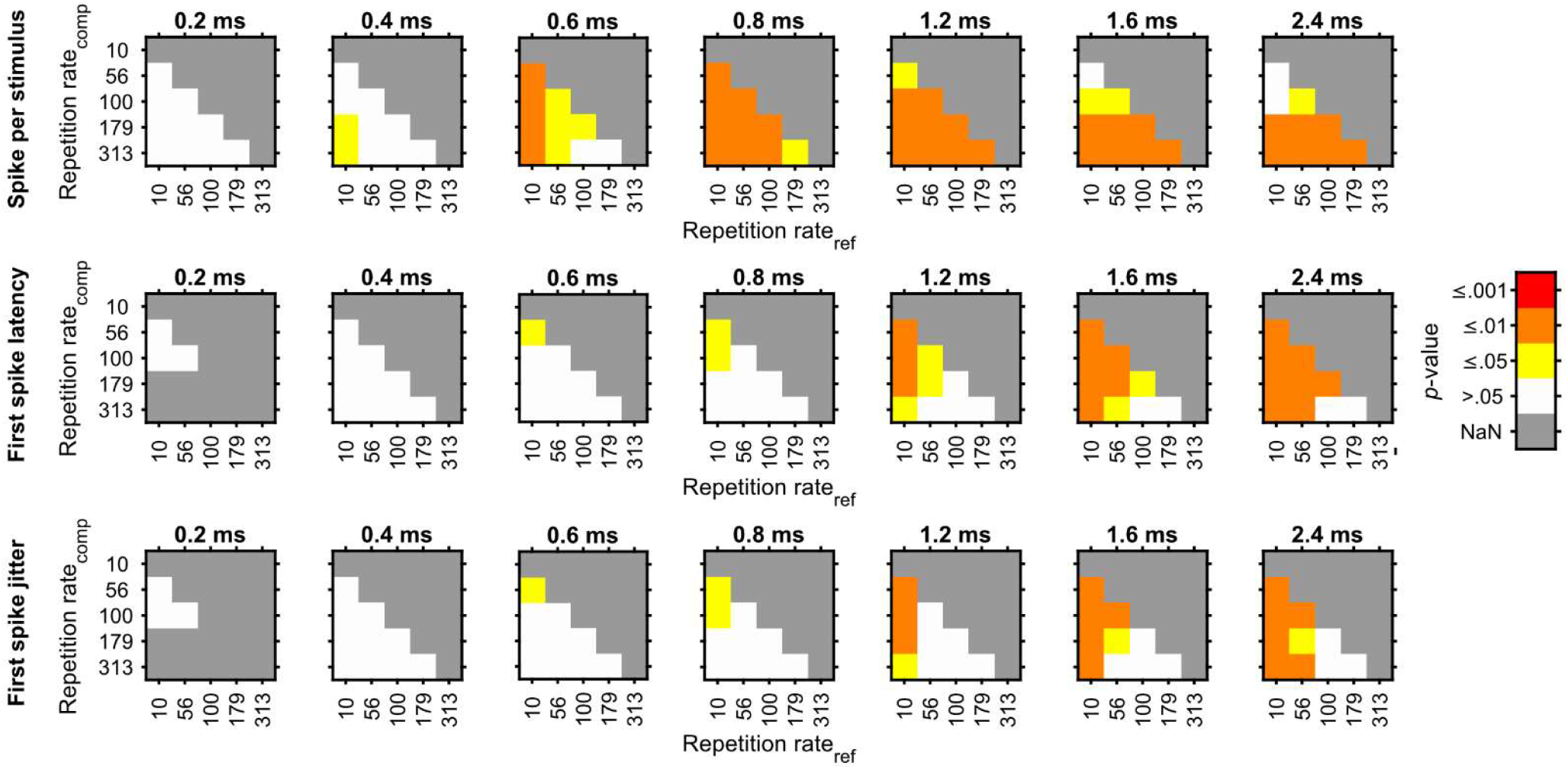
Results of the hypothesis tests performed on the data from figure 5.E-F. to evaluate the effect of the repetition rate at various pulse duration on the number of spikes per stimulus, first spike latency and first spike jitter. Wilcoxon signed rank test on paired samples followed by a Bonferroni correction test. A color code was used to represent the *p*-values.

## Methods

### Animals

Data were obtained from 73 adult (> 12 weeks of age) C57Bl/6 wild-type mice of either sex. For all procedures, animals were placed on a heating pad and body temperature was maintained at 37 °C. All experimental procedures were done in compliance with the German national animal care guidelines and approved by the local animal welfare committee of the University Medical Center Göttingen, as well as the animal welfare office of the state of Lower Saxony, Germany (LAVES).

### Virus purification

The virus purification procedure was performed as previously published (Keppeler et al., 2018) and an extensive description is available in (Huet and Rankovic, 2021). In brief, triple transfection of HEK-293T cells was performed using the pHelper plasmid (TaKaRa, USA), trans-plasmid with PHP.B capsid (Ben Deverman, Viviana Gradinaru, Caltech, USA) and cis-plasmid with Chronos-ES/TS under the control of human synapsin promotor. Cells were regularly tested for mycoplasma contamination. Viral particles were harvested 72 hours after transfection from the medium and 120 h after transfection from cells and the medium; precipitated with 40% polyethylene glycol 8000 (Acros Organics, Germany) in 500mM NaCl for 2 h at 4 °C and then after centrifugation combined with cell pellets for processing. Cell pellets were suspended in 500mM NaCl, 40mM Tris, 2.5mM MgCl2, pH 8, and 100 Uml*−*1 of salt-activated nuclease (Arcticzymes, USA) at 37 °C for 30 min. Afterward, the cell lysates were clarified by centrifugation at 2000 ×g for 10 min and then purified over iodixanol (Optiprep, Axis Shield, Norway) step gradients (15%, 25%, 40%, and 60%) at 350,000 × g for 2.25 h. Viruses were concentrated using Amicon filters (EMD, UFC910024) and formulated in sterile phosphate-buffered saline (PBS) supplemented with 0.001% Pluronic F-68 (Gibco, Germany). Virus titers were measured using AAV titration kit (TaKaRa/Clontech) according to the manufacturer’s instructions by determining the number of DNase Iresistant vg using qPCR (StepOne, Applied Biosystems). The purity of produced viruses was routinely checked by silver staining (Pierce, Germany) after gel electrophoresis (Novex™ 4–12% Tris–Glycine, Thermo Fisher Scientific) according to the manufacturer’s instruction. The presence of viral capsid proteins was positively confirmed in all virus preparations. Viral stocks were kept at *−*80 °C until injection.

### Virus injection

Virus injection was performed as previously published (Keppeler et al., 2018) and an extensive description is available (Huet and Rankovic, 2021). In brief, PHP.B vectors carrying the targeting-optimized Chronos (Chronos-ES/TS) linked to the reporter-protein eYFP and under control of the human synapsin promotor (titer: 3.3 - 8.4 × 10^12^ genome copies/ml) was injected in the left ear at P5-P7. Under general isoflurane anesthesia and analgesia achieved through local application of xylocaine and subcutaneous injection of burprenorphine (0.1 mg/kg) and carprofen (5 mg/kg), the left bulla was approached via a dorsal incision. Injection of 1 – 1.5 μl of virus suspension (mixed with fast green, 1:20) was performed in the scale tympani using laser-pulled (P-2000, Sutter Instrument Inc., USA) quartz capillaries (Science Products, Germany) connected to a pressure micro-injector (100–125 PSI, PLI-100 picoinjector, Harvard Apparatus). After virus application, the tissue above the injection site was repositioned and the wound was sutured. Recovery of the animals was then daily tracked the first week and then weekly until the electrophysiological recordings. If needed, additional application of carprofen (5 mg/kg) could be performed. Animals were kept in a 12-h light/dark cycle, with access to food and water *ad libitum*.

### Stimulation

Stimuli were generated via a custom-made system based on NI-DAQ-Cards (NI PCI-6229; National Instruments; Austin, USA) controlled with custom-written MATLAB scripts (The MathWorks, Inc.; Natick, USA). Acoustic stimuli (sampling rate = 830 kHz) were presented open-field via a loudspeaker (Avisoft Inc., Germany) localized on the left side at ∼15 cm from the pinna. A 0.25-inch microphone and measurement amplifier (D4039; 2610; Brüel & Kjaer GmbH, Naerum, Denmark) were used to calibrate sound pressure levels. Optical stimuli were delivered at the cochlear round window via an optical fiber (200 μm diameter, 0.39 NA, Thorlabs GmbH, Germany) coupled to a blue laser (473 nm, MLLFN-473-100, 100 mW diode pumped solid state [DPSS]; Changchun New Industry Optoelectronics). The maximum radiant flux was measured before every experiment (LaserCheck; Coherent Inc.) and later used for calibration. The round window was exposed as previously described (Bali et al., 2021; Huet and Rankovic, 2021; Keppeler et al., 2018; Mager et al., 2018). Briefly, the round window was exposed by opening the temporal bone ventrally from the *stylomastoid foramen* and the exact location of the round window was determined by visually following the stapedial artery.

### Auditory brainstem recordings (ABR)

Animals were anesthetized via intraperitoneal injection of a mixture of xylazine (5 mg/kg) and urethane (1.32 mg/kg) and appropriate analgesia was achieved by sub-cutaneous injection of buprenorphine (0.1 mg/kg, repeated every 4 hours) and carprofen (5 mg/kg). Depth of anesthesia was monitored regularly by the absence of reflexes (hind limb withdrawal) and adjusted accordingly. ABRs were recorded using sub-dermal needle electrodes inserted underneath the pinna, on the vertex, on the back near the contralateral leg and signals were amplified using a custom-made differential amplifier, sampled at a rate of 50 kHz, filtered (second order Butterworth filter, 300 – 3000 Hz) and averaged across 1000 iterations. The first ABR wave, reflecting the synchronous activation of the SGNs, was detected semi-automatically with a custom-written MATLAB script. Thresholds were visually defined as the lowest intensity that elicited a reproducible response in the recorded traces.

### Juxtacellular recordings of SGNs and AVCN neurons

If positive acoustic or optogenetic ABRs were recorded, the mice were prepared for subsequent juxtacellular recordings of the SGN and AVCN neurons. Briefly, a tracheotomy and intubation were performed before mounting the animal in a custom-designed stereotactic head holder. Pinnae were removed, scalp reflected, portions of the lateral interparietal and the left occipital bone removed, and a partial cerebellar aspiration was performed to expose the left semi-circular canal. A reference electrode was placed on the contralateral muscles behind the right ear. A borosilicate sharp micropipette (∼ 50 mΩ, 3M NaCl) was navigated from a reference point at the surface of the semi-circular canal to a stereotaxic position from where SGNs and AVCN neurons recordings could be obtained (LN Mini 55 micro-manipulator, Luigs & Neumann, Germany). Using the step function from the micromanipulator (3 mm/s, 1.5 μm steps), juxtacellular recordings from SGNs or AVCN neurons were obtained. At the end of the procedure, the relative position of the auditory meatus was visually controlled by exposing it. Signals were amplified (ELC-03XS, NPI electronic, Germany) and digitized at a sampling rate of 50 Hz by the same NI-DAQ-Cards and custom-written MATLAB scripts used for stimulus generation. Spikes were detected offline based on their threshold (manually determined) using custom-made MATLAB scripts.

Differentiation between SGN and AVCN neurons was performed on the sole basis of 3 criteria. Units were classified as SGNs if their depth relative to the cerebellum surface ≥ 1000 μm, their first spike latency ≤ 3 ms in response to 0.8 - 1.6 ms light pulse presented at maximum radiant flux / 80 dB SPL_PE_ acoustic click, and if their action potentials were monophasic and positive.

### Stimulation strength

The stimulation strength was defined acoustically as the sound pressure level (SPL) in dB SPL_PE_ (PE: peak equivalent, Cody & Russell, 1987; Kiang *et al*, 1965) and optogenetically as the pulse duration. Previous measures have shown that the oABR amplitude increases with the pulse duration, thus suggesting an increased firing synchronization between SGNs (**figure S1.C**, (Keppeler et al., 2018)) with that parameter. Evaluation of the stimulus strength was either achieved by single stimulus (presented at 10 Hz, 80 presentations) or stimulus trains (100 – 1000 Hz repetition rate, 20 iterations, 400 ms light/sound, 100 ms dark/silence). To optimize acquisition time and enable pairwise comparisons, 2 s stimulation containing 4 blocks of 500 ms were built (400 ms). Acoustically, pairwise acquisitions of the 4 different repetitions rates (10, 100, 316 and 1000 Hz) were performed. Following successful acquisition at 80 dB SPL_PE_, acquisitions were repeated at either 60 and 70 dB SPL_PE_ in a random manner. Only units acquired at the 3 sound intensities were included. Optogenetically, pairwise acquisitions of 4 different light pulse durations (0.4, 0.8, 1.6 and 3.2 ms) were performed at either 10, 100, 316 or 1000 Hz (if the pulse duration exceeded the stimulation period, no acquisition was done for that condition).

The number of spikes evoked per stimulus was counted from the 10 ms following each stimulus presentation. The first spike following each stimulus was detected and its average latency and latency jitter (i.e. standard deviation of the latency) were computed. The adaptation ratio was computed as the ratio between the discharge rate during the first 50 ms of the train and the whole duration of the stimulation train.

### Population response

To estimate the representation of the stimulus across a population of SGNs, we computed the number of spikes per stimulus, first spike latency and first spike latency jitter across recorded SGNs by computationally mimicking a concomitant recording of the SGN population in response to one stimulus. This was achieved by pooling together the response of all recorded SGNs to one randomly selected iteration of the stimulus per SGN, computing the metrics of interest and repeating this operation 50 times (e.g. bootstraps).

### Intensity

Intensity analysis was performed varying pairwise light pulse duration (0.4, 0.8, 1.6 and 3.2 ms) and radiant flux of a given stimulus at 100 Hz presented above. Stimulus amplitude was related to the threshold of auditory brainstem responses (see (Bali et al., 2021)). Radiant flux (light pulse amplitude) was expressed in dB_oABR threshold (mW)_ = 10 × log10 (A/A0) where A is the presented radiant flux and A0 the radiant flux at oABR threshold. Stimuli were presented first at maximum laser output and repeated at as many other intensities as the unit could be held to cover the full dynamic range (i.e. from threshold to saturation). Only units for which at least 4 intensities were measured were included. We previously showed that SGN firing at the threshold is characterized by a strong adaptation (Bali et al., 2021), therefore rate-level functions (RLF) and subsequent analysis were only computed from the plateau response (between 100 and 400 ms). The spontaneous activity of each neuron was measured from the lowest sub-threshold intensity at 0.4 ms and RLF were classified as responding if their discharge rate at the highest intensity was 10 spikes/s above the spontaneous discharge rate. Responding RLFs were classified as saturating if the slope measured between the discharge rate at the two highest measured intensities was ≤ 3 spikes/dB, else they were classified as non-saturating. Per RLF, if sub-threshold intensities were measured, we computed interpolated d’-level function (with the spontaneous activity as a reference, linear interpolation with a 0.2 dB step) from which we defined the threshold as the lowest intensity with a d’ ≥ 1 (Bali et al., 2021; Huet et al., 2018; Macmillan and Creelman, 2004). Saturated RLFs were fitted with a sigmoidal equation and the dynamic range (i.e. the difference in intensity) and light-driven rate (i.e. the difference in discharge rate) were measured between the threshold and the intensity leading to 95% of the maximum discharge rate. Dynamic range and light-driven rate from unsaturated RLFs were measured between the threshold and the maximum tested intensity.

### Forward masking protocol

Recovery from the optogenetically induced firing refractoriness was measured by a so-called forward masking protocol. Ten light pulses of 1.6 ms presented at 313 Hz were used as a masker to induce firing adaptation and the recovery was measured at different time intervals by a single light pulse (i.e. the probe) of 1.6 ms. A total of 20-time intervals ranging between 4 and 180 ms were tested and a 200 ms dark time was applied between each measure. Each interval was tested 20 times. Stimuli were presented first at maximum laser output and repeated at as many other intensities as the unit could be held. The success of the masker to fully adapt the firing was evaluated from the discharge rate and the adaptation ratio of the firing in response to the masker. Next, the spike probability (i.e. the probability of the probe to evoked spikes) per interval was measured and normalized to the spike probability in response to the longest interval. Recovery curves were fitted by a mathematical model (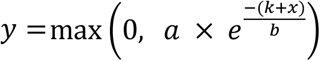 to define the time to absolute recovery (i.e. time required to recover any firing, - k) and relative recovery (i.e. time required to recover 95 % of the spike probability, - k + (3 x b)). Additionally, the spontaneous discharge rate of the SGNs was measured in the last 180 ms of the 200 ms dark time following each probe.

### Semi-stochastic stimulus

A semi-stochastic stimulus was designed for pairwise testing of 35 combinations of repetition rates (10, 56, 100, 179 and 313 Hz) and light pulse durations (0.2, 0.4, 0.6, 0.8, 1.2, 1.6, 2.4 ms). Each combination was randomly tested 10 times per stimulus iteration and the presentation order was randomized for each of the 20 iterations, resulting in 200 presentations per combination. Each iteration started with a masker composed of 10 light pulses of 2.4 ms presented at 100 Hz and finished with 200 ms of dark. Additionally, a forward masking protocol testing for the intervals of 15 and 80 ms was included. Recordings were made at maximum radiant flux. Similarly, an acoustic stimulus was designed using acoustic click (100 μs) and testing for 5 repetition rates (10, 56, 100, 179 and 313 Hz). Recordings were made at 60, 70 and 80 dB SPL_PE_. Raster plots were built from a 3 ms window following each stimulus which was defined by first computing a spike-triggered average. For each condition, the number of spikes per stimulus, first spike latency and first spike latency was computed as described above.

### Cochlear histology

Following each optogenetic injection, cochleae from both side were prepared as previously described (Bali et al., 2021; Keppeler et al., 2018; Mager et al., 2018). Briefly, samples were fixed for an hour in 4% paraformaldehyde (PFA), decalcified for 3-4 days in 0.12 M ethylenediaminetetraacetic acid (EDTA), dehydrated in 25% sucrose solution for 24 hours and cryosectioned (mid-modiolar cryosections, 25 μm tick). Immunofluorescence staining were done in 16% goat serum dilution buffer (16% normal goat serum, 450 mM NaCl, 0.6% Triton X-100, 20 mM phosphate buffer, pH 7.4). The following primary antibodies were used at 4 °C overnight: chicken anti-GFP (1:500, ab13970 Abcam, USA) and guinea pig anti-parvalbumin (1:300, 195004 Synaptic Systems, Germany); and secondary antibodies for one hour in room temperature: goat anti-chicken 488 IgG (1:200, A-11039 Thermo Fisher Scientific, USA) and goat anti-guinea pig 568 IgG (1:200, A-1107 Thermo Fisher Scientific, USA). Finally, slices were mounted in Mowiol 4-88 (Carl Roth, Germany). Samples were imaged with a LSM 510 Zeiss Confocal Microscope (Zeiss, Germany) mounted with a 40x air objective. An image per cochlear turn (apex, middle and base) was taken from the modiolus (i.e. the bony structure containing the SGN somas). Image analysis was performed by a custom-written MATLAB script modified from (Huet et al., 2021). Briefly, SGN somas and modiolus area were manually detected using a touch screen from the parvalbumin images. Next, individual somas were automatically segmented using Otsu’s threshold method from every Z-stack and a mask corresponding to the given SGN was defined for the Z-stack for which the mask was fulfilling the criteria of size (area and diameter) and circularity. In case the segmentation was not right, the segmentation of the given SGN soma was performed manually. Next, the median GFP brightness of each SGNs was measured and its distribution was fitted with a Gaussian mixture model with up to 3 components. A threshold, above which SGNs somas were considered as transduced, was defined as average + 3 x standard deviation of the Gaussian distribution with the lowest mean.

### Analysis

Data were analyzed using Matlab (MathWorks). All averages in text and figures were expressed as mean ± 95% confidence interval. Normality was tested by a Jarque-Bera test. For statistical comparison between 2 independent groups, two-sample t-test (parametric) or a Wilcoxon rank sum test (non-parametric) tests were used. For statistical comparison between more than 2 independent groups, ANOVA (parametric) or Kruskal-Wallis test (non-parametric) were used followed by a Tukey’s Honest Significant Difference procedure. For statistical comparison between paired data, a Wilcoxon signed rank test on paired samples followed by a Bonferroni correction of the p-values were used.

## Acknowledgments

We warmly thank Marcus Jeschke, Thomas Mager and Lina Maria Jaime Tòbon for critical discussions influencing the design of the study and the interpretation of the results. We thank Anupriya Thirumalai for assisting with the oABR and histological analysis. We thank Burak Bali for teaching the single unit recordings of the SGNs. We thank Bettina Wolf for her general support and critical insights. We thank Vladan Rankovic for virus preparation and virus injection. We thank Daniela Gerke for expert help with virus and immunolabeled midmodiolar cochlear cryosection preparation, Christiane Senger-Freitag and Sandra Gerke for expert technical support, Gerhard Hoch for engineering support, and Patricia Räke-Kügler for excellent administrative support. We also thank Ben Deverman and Viviana Gradinaru for providing the PHP.B construct used in this study. A.M. and A.H. thank the Hertha Sponer College for providing an intellectually stimulating environment where to exchange about science.

This work was funded by the European Research Council through the Advanced Grant ‘OptoHear” to T.M. under the European Union’s Horizon 2020 Research and Innovation program (grant agreement No. 670759), the Fraunhofer and Max-Planck cooperation program (NeurOpto grant) to T.M., the German Research Foundation through the Priority Program 1926 “Next generation optogenetics” to A.H. and T.M. and was further supported by the German Research Foundation through the Cluster of Excellence (EXC2067) Multiscale Bioimaging to T.M. and A.H., as well as the Leibniz Program to T.M. A.M. is recipient of a scholarship of the Göttingen Promotionskolleg für Medizinstudierende, funded by the Jacob-Henle-Programm or Else-Kröner-Fresenius-Stiftung (Promotionskolleg für Epigenomik und Genomdynamik, 2017_Promotionskolleg.04). In addition, this research was supported by Fondation Pour l’Audition (FPA RD-2020-10) to TM.

## Author contributions

A.M, T.M. and A.H. designed the study. A.M. performed oABR recordings, single-unit recordings of SGNs and AVCN neurons and SGNs histology. A.M. and A.H analyzed the data and A.H. performed computation. A.M., T.M. and A.H. prepared the manuscript.

## Notes

### Competing Interest Statement

Tobias Moser is a co-founder and CEO of OptoGenTech company. The other authors declare no conflict of interests.

### Summary of Updates

In this version of the manuscript, we strengthened the impact of our result on the design of the future optical cochlear implant.

